# Enterovirus D68 capsid formation and stability requires acidic compartments

**DOI:** 10.1101/2023.06.12.544695

**Authors:** Ganna Galitska, Alagie Jassey, Michael A. Wagner, Noah Pollack, William T. Jackson

**Affiliations:** Department of Microbiology and Immunology, University of Maryland School of Medicine, 685 W. Baltimore St, Baltimore, MD 21201, USA

**Keywords:** picornaviruses, autophagy, enterovirus D68, poliovirus, cellular acidification, virion assembly, virus maturation, autophagosomes, replication organelles (ROs), multi-vesicular bodies (MVB), “transition point”

## Abstract

Enterovirus D68 (EV-D68), a picornavirus traditionally associated with respiratory infections, has recently been linked to a polio-like paralytic condition known as acute flaccid myelitis (AFM). EV-D68 is understudied, and much of the field’s understanding of this virus is based on studies of poliovirus. For poliovirus, we previously showed that low pH promotes virus capsid maturation, but here we show that, for EV-D68, inhibition of compartment acidification during a specific window of infection causes a defect in capsid formation and maintenance. These phenotypes are accompanied by radical changes in the infected cell, with viral replication organelles clustering in a tight juxtanuclear grouping. Organelle acidification is critical during a narrow window from 3-4hpi, which we have termed the “transition point,” separating translation and peak RNA replication from capsid formation, maturation and egress. Our findings highlight that acidification is crucial only when vesicles convert from RNA factories to virion crucibles.

**Importance:** The respiratory picornavirus enterovirus D68 is a causative agent of Acute Flaccid Myelitis, a childhood paralysis disease identified in the last decade. Poliovirus, another picornavirus associated with paralytic disease, is a fecal-oral virus which survives acidic environments when passing from host-to-host. Here we follow up on our previous work showing a requirement for acidic intracellular compartments for maturation cleavage of poliovirus particles. Enterovirus D68 requires acidic vesicles for an earlier step, assembly and maintenance of viral particles themselves. These data have strong implications for the use of acidification blocking treatments to combat enterovirus diseases.

## Introduction

Poliomyelitis, one of the most feared diseases of the twentieth century, is now limited to regional outbreaks of unvaccinated patients, and declarations of poliovirus (PV) eradication are perpetually imminent.(1) With the public health relevance of PV lessened, the related picornavirus Enterovirus D68 (EV-D68) has come into the spotlight, due to its association with the polio-like neurological disease Acute Flaccid Myelitis.(2) Although EV-D68 was first described over a half century ago, most hypotheses regarding the interactions of EV-D68 with the host cell, and its viral cycle, rely on assumptions from work done on poliovirus and a few other picornaviruses.

Despite being defined as an “enterovirus,” EV-D68 is primarily associated with respiratory symptoms.(3) Infections were sporadically reported in the US, Asia, and Europe until a large outbreak took place in 2014 in North America, which was particularly dangerous to children with a history of asthma or reactive airway disease.(4–7) With the exception of the coronavirus pandemic year of 2020, EV-D68 cases have since been reported with biennial and seasonal patterns, and from 2014-2018 these coincided with the occurrence of acute flaccid myelitis (AFM) cases in children.(8–10) AFM manifests as a rare polio-like neurological condition characterized by acute onset of flaccid limb weakness, lesions in gray matter, or, eventually, paralysis.(11–14) EV-D68 is one of a few pathogens considered to be a likely cause of AFM, yet little is known about the relative life cycles of poliovirus and enterovirus D68.(2, 15) A better understanding of their comparative life cycles will be crucial to understanding the differences and similarities between poliomyelitis and AFM. In general there is a need for a specific understanding of EV-D68 infections, as no vaccines or treatments are currently available for the latter.(16)

As a member of the picornavirus family, EV-D68 contains a single-strand positive RNA genome and shares biological features with other enteroviruses, including coxsackievirus and poliovirus.(17) All picornaviruses co-opt cellular processes in hosts to facilitate the replication of their genomes and spread to neighboring cells. In such a manner, they target autophagy (also often referred to as macroautophagy), which is one of the most crucial and conservative stress response pathways that allow cells to recycle cellular components by sequestering various molecules and organelles in double-membraned vesicles (DMVs), also known as autophagosomes, which fuse with lysosomes, resulting in degradation of the autophagic cargo.(18–20)

Autophagy is an “always on” process, but levels of autophagic activity are regulated by mTOR and Akt, which signal to the pathway-specific complex consisting of ULK1, RB1CC1, ATG13, and ATG101.(21–23) The kinase member of that complex, ULK1, signals to the Beclin1/ATG14/PIK3C3 complex, which in turn induces lipidation and membrane insertion of the LC3 protein, the earliest protein marker of autophagosomes.(24–29) The SQSTM1 cargo adaptor binds to both LC3 and cellular cargo, sequestering the cargo in autophagosomes. SQSTM1 is degraded with cargo, making it an excellent monitor for active autophagic degradation.(28–30) Autophagosomes fuse with endosomes to acidify into amphisomes; this acidification facilitates fusion with lysosomes for cargo degradation.(31)

Many picornaviruses utilize the autophagic machinery to reorganize the membrane surfaces into replication sites and use them as a source of vesicles for maturation and exit from cells.(18, 32–35) The pro-viral role of autophagy has been well documented for poliovirus: the virus triggers the formation of complex membrane structures known as membrane replication complexes, MRCs, also known as replication organelles (ROs,) which act as the physical site of viral RNA (vRNA) replication. After the peak of RNA replication, the virus usurps the autophagy machinery to convert ROs to autophagosome-like DMVs, which mature into acidic amphisomes. PV depends on vesicle acidification for its virion maturation, and mature virions use an exit pathway we previously termed autophagosomal exit without lysis (AWOL).(18, 36–39) We and other groups have demonstrated that multiple picornaviruses, including EV-D68, initiate the activation of autophagy pathways while cleaving host SNARE protein SNAP29 to prevent autophagosome/lysosome fusion. In addition, both CVB3, EV-D68, and poliovirus have been shown to cleave SQSTM1, thus inhibiting autophagic cargo degradation via selective autophagy.(40, 41)

PV has been shown to induce autophagy independent of canonical ULK1 signaling, but this has not been tested for EV-D68.(41) The role of autophagosome-like vesicles, and the importance of organelle acidification to the formation of the cellular membrane replication complexes supporting EV-D68 replication, virion synthesis, assembly, virus maturation, and its exit from the host cell has been generally assumed by our group and others to be similar to that of poliovirus. Here we investigate the fundamental differences in life cycles between the PV and EV-D68 and how these viruses are able to exploit and evade host autophagy. We show that EV-D68 growth is delayed under acidification inhibitors, and that entry is blocked. We also show that vRNA levels drop when acidification inhibitors are present, and that, depending on the timing of inhibitor addition, capsid formation and/or stability are severely affected, and an unusual clustering of the ROs is observed. Our findings on the role of cellular compartment acidification, and its contribution to successful generation of virus progeny, provides insights that will pave the path to understanding unique aspects of EV-D68 infection.

## Results

### EV-D68 and PV display similarities in the kinetics of virus production and interaction with the canonical signaling pathway

Given that picornavirus replication cycles have many common characteristics, we sought to compare growth between poliovirus and enterovirus D68 using H1HeLa cells, which serve as an established model for studying picornaviruses.(42) We first assessed virus production by measuring viral replication kinetics for intracellular and extracellular virus titers at various times post-infection. Both viruses display similar kinetics of viral production with a rapid exponential increase in the production of infectious virus particles relatively early in the life cycle, between 2 to 4 hours post-infection (hpi) (Figure 1A). Infected cells do not undergo cell lysis until later time points (4.5-6 hpi, data not shown), suggesting that non-lytic release of infectious virus particles is possible for both viruses. We then analyzed the impact of EV-D68 infection on autophagy markers in infected cells and observed that EVD68, like PV, induces both gradual accumulation of LC3BII over the course of infection and cleavage of SQSTM1. We conclude that, like PV, EV-D68 induces markers of autophagy but blocks autophagic flux, thus preventing degradative autophagy (Figure 1B).(43)

**Figure 1.**
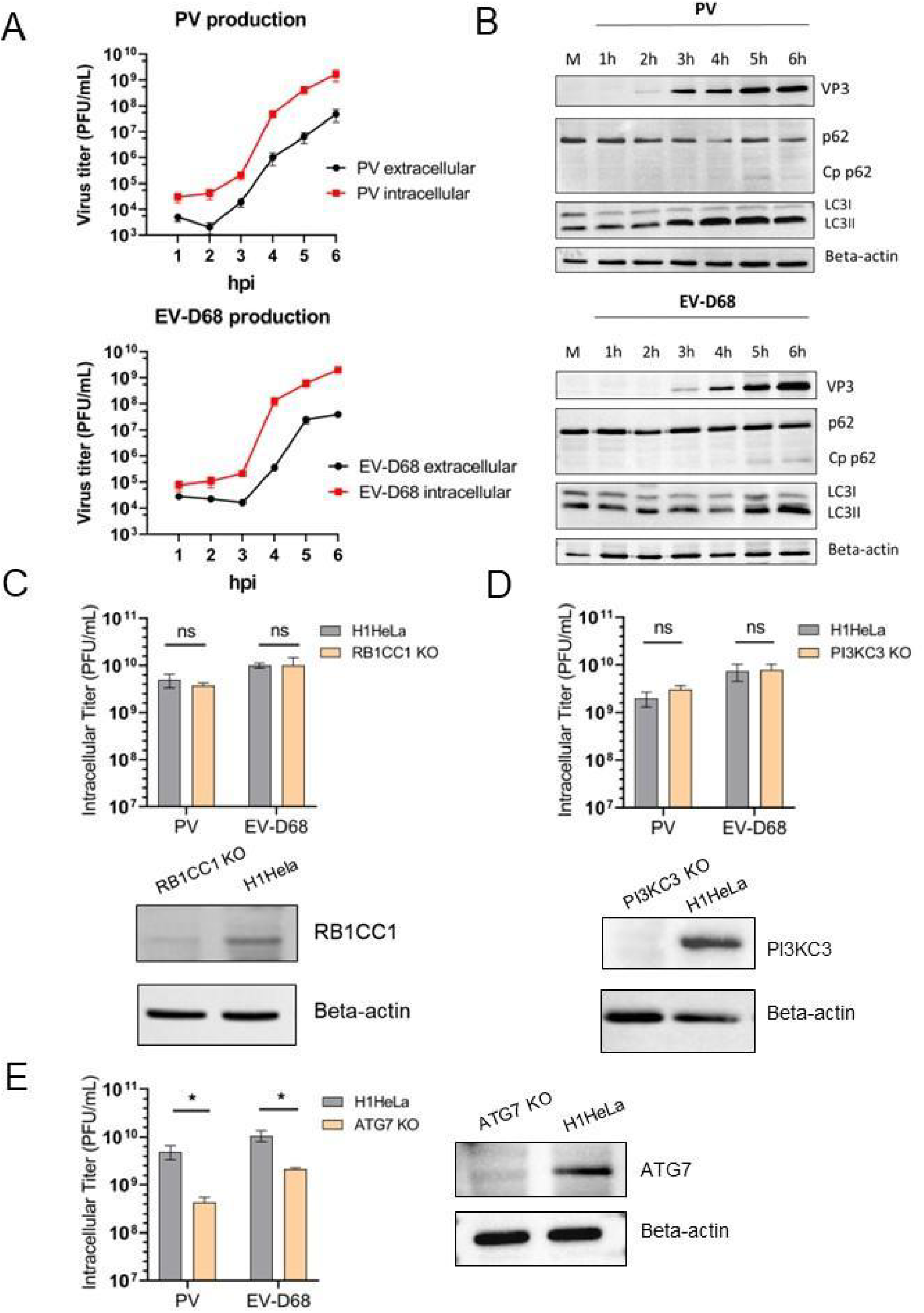
EV-D68 and PV display similarities in virus production kinetics and interaction with canonical autophagy players. (A) Titers of both cell-associated and extracellular PV and EV-D68 from a time course of infection. H1HeLa cells were infected at an MOI of 20. Intracellular and extracellular virus samples were collected at each hour post-infection. Viral titers were analyzed by plaque assay. (B) Impact of enterovirus D68 and poliovirus on host autophagy markers. H1HeLa cells were either untreated/mock (M) or infected with PV or EV-D68 at an MOI of 20 for 6 h, and samples were collected every hour during infection. Samples were subjected to western blot analysis for traditional autophagy markers: LC3B, p62(SQSTM1), and its Cp (cleavage product). Viral marker: VP3 (virus structural capsid protein 3). Beta-actin served as a loading control. (C-E) Intracellular EV-D68 and PV titers produced in parental H1HeLa vs. KO cell lines. H1HeLa WT or KOs were infected with PV or EV-D68 at an MOI of 20 for 6 h. Viral titers were analyzed by plaque assay. KO cell lines represented: RB1CC1 KO (C), PI3K3 KO (D), and ATG7 KO(E), respectively. Unpaired student’s t-test was used for the statistical analysis (***=p < 0.001; **= p< 0.01; *= p ≤ 0.05; ns=not significant). The efficiency of KOs was assessed by Western Blot and the parental H1HeLa cell line (WT) was used as a control.

To investigate the role of autophagy in the life cycle of EV-D68, we generated individual CRISPR/CAS9 H1HeLa cell knock-outs (KOs) in members of the three major regulatory complexes of the canonical autophagic signaling pathway: RB1CC1 (ULK complex); PI3KC3 (Beclin1 complex); and ATG7 (LC3 lipidation complex). Both confirmed KOs (Figure 1C-E) and parental H1HeLa cells were infected with either PV or EV-D68 (MOI=20) and the cell-associated virus titers were measured at 6 hpi (Figure 1C-E). Our results demonstrate that EV-D68, similarly to PV, does not rely on canonical autophagy signaling (Figure 1C,D) for successful virus production. However, both viruses require functional ATG7 to generate infectious progeny (Figure 1E). Therefore, our data show that both PV and EV-D68 interact similarly with autophagic pathways and require intact ATG7 protein that drives the stages of classical degradative autophagy via LC3 lipidation for the formation of double-membrane vesicles; however, this process occurs independently of canonical autophagic upstream signaling.

### Poliovirus and EV-D68 respond differently to acidification inhibitors

While both viruses block autophagic flux, we wanted to understand how the EV-D68 virus life cycle benefits from this process. Autophagosome maturation can be manipulated by inhibitors of vesicle acidification, including ammonium chloride (NH_4_Cl), the vacuolar inhibitor of H+-ATPases Bafilomycin A1 (Baf A1), and chloroquine (CQ), which successfully inhibit amphisome-lysosome fusion, as well as subsequent cargo degradation. These inhibitors are used in autophagy research as interchangeable, although their mechanisms of action are somewhat different.(44, 45) While the weak base ammonium chloride immediately increases intralysosomal pH via proton trapping, Bafilomycin A1 inhibits autophagosome-lysosome fusion by specifically blocking H(+)-ATPase pump activity.(46–48) Chloroquine, as a lysotropic weak base, in the monoprotonated form diffuses rapidly and spontaneously across the membranes of various cell organelles, including lysosomes, where it becomes deprotonated and becomes trapped. Protonated chloroquine then changes the lysosomal pH, inhibiting autophagic degradation in the lysosomes.(49) Moreover, CQ raises pH in other acidic vesicles, including endosomes and Golgi vesicles, but has also been proven to cause multiple effects, including dysfunction of various enzymes and organelles such as mitochondria, and even altering immune processes, such as lysosome-mediated antigen presentation.(50–55)

To further investigate the importance of blocking autophagic flux on EV-D68 production, we treated cells with these acidification inhibitors. Previously, our lab reported the impact of acidification inhibitors on PV titer, with a specific requirement for low pH conditions to promote maturation of infectious PV particles within autophagosomes.(37) Western Blots of untreated H1HeLa vs. cells treated with acidification inhibitors have been performed to monitor the accumulation of LC3B-II as an indicator of blocked autophagy (Figure 2A, top right; Figure S1A-B). Upon treatment with these acidification inhibitors, intracellular titers of EV-D68 remain unaffected, in contrast to PV, the titer of which was significantly reduced upon treatment (Figure 2A-C).

**Figure 2.**
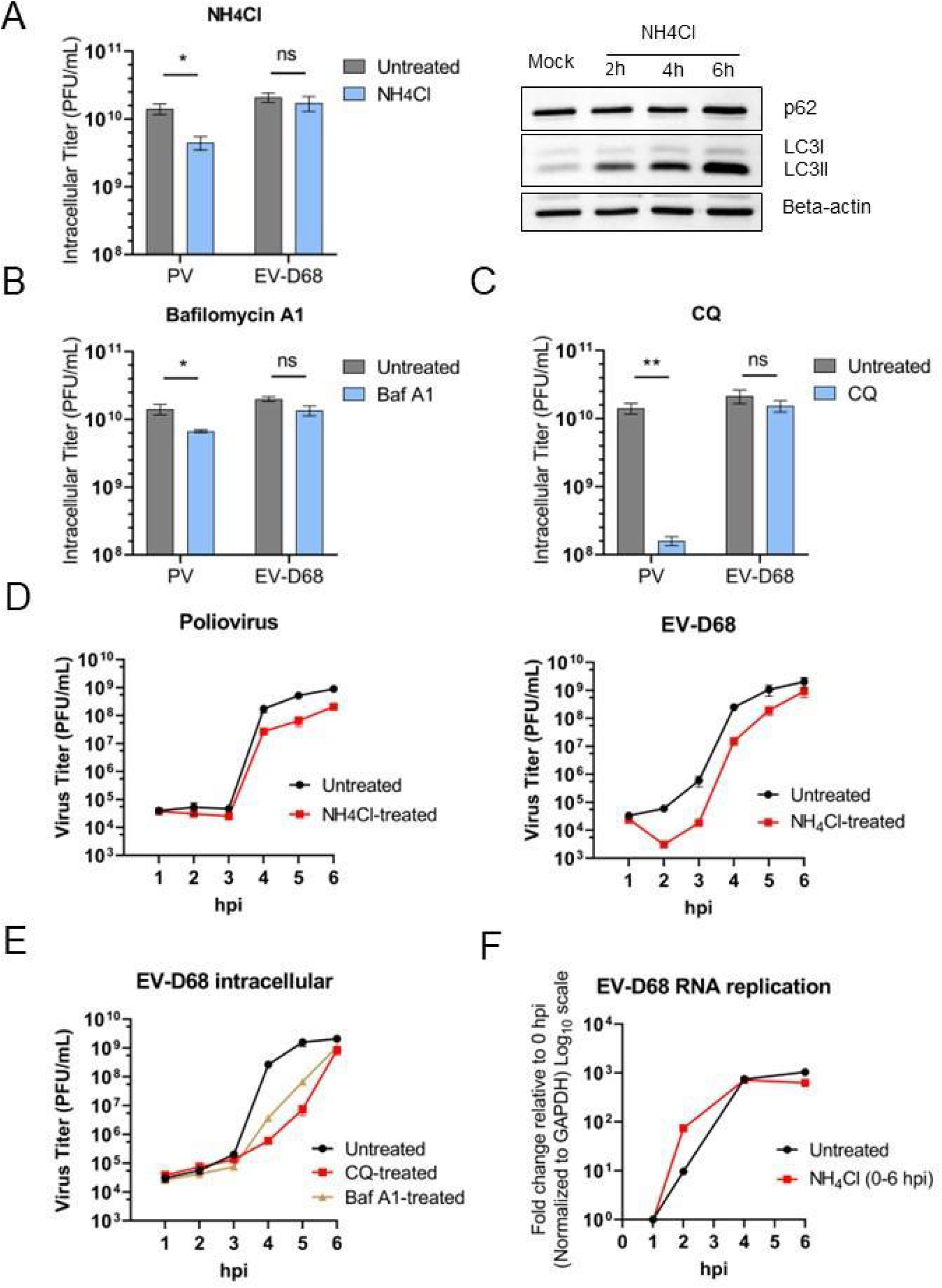
Poliovirus and EV-D68 respond differently to acidification inhibitors with major differences in virus production. (A) Left panel: titers of cell-associated PV and EV-D68 upon treatment with ammonium chloride (NH_4_Cl, 20mM). H1HeLa were infected with PV or EV-D68 at an MOI of 20; after 30 min of virus adsorption at 37°C (indicated as 0 hpi time point), cells were washed twice to remove the uninternalized virus particles, then left either untreated/mock (M) or treated with ammonium 20mM in a complete medium throughout the infection. Cells were collected at 6 hpi. Right panel: autophagy markers SQSTM1 and LC3B in mock samples of H1HeLas, untreated or treated with ammonium chloride (NH_4_Cl, 20mM) for 2h, 4h, or 6h, were analyzed by Western blot. Beta-actin served as a loading control. The gradual accumulation of LC3B served as an indicator of blocked autophagy. (B) Titers of cell-associated PV and EV-D68 upon treatment with Bafilomycin A1 (Baf A1, 0.1uM). H1HeLa were infected with PV or EV-D68 at an MOI of 20; after 30 min of virus adsorption at 37°C (0 hpi), cells were washed twice to remove the initial virus input, left untreated, or treated with Baf1A (0.1uM) in a complete medium throughout the infection. Cells were collected at 6 hpi. (C) Titers of cell-associated PV and EV-D68 upon treatment with chloroquine (CQ; 100uM). H1HeLa were infected with PV or EV-D68 at an MOI of 20; after 30 min of virus adsorption at 37°C (0 hpi), cells were washed twice to remove the initial virus input, left untreated, or treated with chloroquine (CQ; 100uM) in a complete medium throughout the infection. Cells were collected at 6 hpi. (D) Titers of cell-associated EV-D68 and PV from a time course of infection: upon treatment with ammonium chloride vs untreated infection. H1HeLa cells were infected at an MOI of 20 for 30 min at 37°C. After 30 min, at the “0 hpi” time point, the cells were washed twice to remove the residual virus, and either treated with ammonium chloride (NH_4_Cl, 20mM) or left untreated. Intracellular samples for plaque assay were collected at each hour. (E) Titers of cell-associated EV-D68 from a time course of infection: upon treatment with either Baf A1 or CQ between 0 hpi to 6 hpi vs untreated infection. Intracellular virus samples were collected at each hour. Viral titers were analyzed by plaque assay. (F) Effect of ammonium chloride treatment on viral RNA synthesis assessed by qRT-PCR. Cells were infected with EV-D68 at an MOI of 20 for 30 min at 37°C, then the residual virus was washed away, followed by either adding normal media (untreated infection, EV-D68; -NH_4_Cl) or media containing 20mM of ammonium chloride (EV-D68; +NH_4_Cl). Cells were then collected at the indicated time points after infection, and total RNA was extracted and subjected to qRT-PCR RNA analysis following cDNA synthesis. The level of GAPDH mRNA was used as an internal control. Unpaired student’s t-test was used for statistical analyses (***=p < 0.001; **= p< 0.01; *= p ≤ 0.05; ns=not significant).

To elucidate the impact of acidification inhibitors on EV-D68 and PV, we titered virus of each time point of the infection, to reveal timepoint-specific differences in viral production between PV and EV-D68 (Figure 2D). As expected, in agreement with previously reported data, poliovirus titers decreased (beginning at 3hpi) upon treatment with ammonium chloride, due to a lack of vesicular acidification that PV relies on for viral particle maturation.(37) As previously reported, upon continuous treatment with ammonium chloride, PV RNA replication is not affected (Figure S1C).

In contrast, our results show that treatment with ammonium chloride results in a brief (approx. 1h) delay in EV-D68 production, which then recovers at the late stages of infection, ultimately reaching the same levels as untreated control infection. A similar EV-D68 production delay was also observed upon treatment with CQ and Bafilomycin A (Figure 2E). This delay in virus production is not due to direct inhibition of vRNA replication upon treatment with ammonium chloride (Figure 2F).

### A delay in EV-D68 production is associated with a delay in viral entry/fusion

To determine if the observed delay in EV-D68 production is due to a delay in viral entry, we performed a synchronized entry assay, as previously described.(56) We tested acidification inhibitor treatments for impact on viral entry/fusion (Figure 3A-B), cell-free virus particles (virus inactivation assay), and viral attachment (Figure S2A-B). The results demonstrate that neither of the acidification inhibitors neutralize the viral particles via viral inactivation assay (Figure S2A), nor has an impact on EV-D68 or PV attachment (Figure S2B). However, PV and EV-D68 differ in their response to acidification inhibitors during entry and fusion: both PV and EV-D68 entry process is strongly inhibited both by CQ and ammonium chloride (Figure 3B). The entry of neither virus was affected by Bafilomycin A1 (Figure 3B), in agreement with what has been previously reported for poliovirus.(57) This suggests that an acidified entry vesicle, but not the activity of Bafilomycin-targeted proton pumps, is required for viral entry.

**Figure 3.**
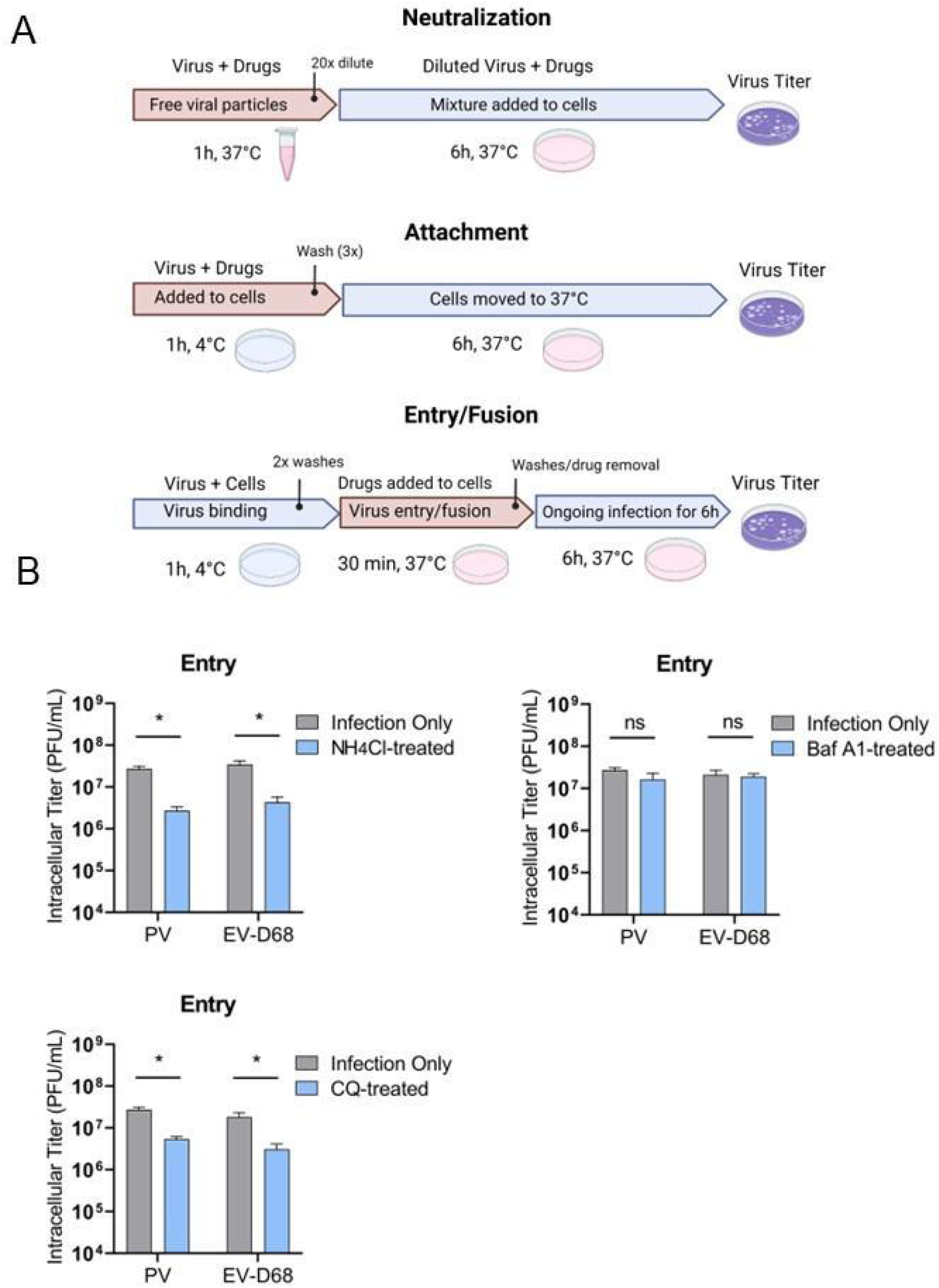
Delay in EV-D68 production is associated with a delay in viral entry/fusion. (A) A schematic of the assays employed to investigate the effect of acidification inhibitors on the early steps of viral cycles. Except for the virucidal neutralization assay, H1HeLa cells were pre-chilled for 30 min on ice before each virus infection. All infections with PV or EV-D68 were performed at 0.1 MOI. Cells were washed twice with PBS (1X) between each incubation period to remove the residual virus after binding or before substituting drug-containing media with normal DMEM. End time points of infections were collected at 6 hpi. All intracellular titers were analyzed by plaque assay. Unpaired student’s t-test was used for the statistical analysis (***= p< 0.001; **= p< 0.01; *= p ≤ 0.05; ns=not significant). (B) Effect of acidification inhibitors on viral entry/fusion. H1HeLa were pre-chilled before the infection of PV or EV-D68 at MOI 0.1. The virus binding/attachment step was allowed to run for 1h at 4°C. Then the residual virus was washed at least twice and the media containing drugs at the defined concentrations (NH_4_Cl 20mM, Bafilomycin A 0.1uM, CQ 100uM) were added to cells and immediately moved to 37°C for 30 min. After 30 min, cells were washed and the media without the drug was added. The infection was allowed to proceed for 6h and then the cells were collected.

### EV-D68 capsid production and virion maturation do not depend on cellular acidification, but the production of several non-structural proteins does

Since EV-D68 viral RNA levels are not affected by continuous ammonium chloride added at 0 hpi (Figure 2F), we investigated the impact of acidification inhibitors on processes of viral capsid production and virion maturation. Assembly of viral capsid proteins is thought to occur as nascent RNA genomes are formed. Capsid protomers, the smallest units of the capsid complex, form into pentamers, and twelve 14S pentamers assemble into the empty capsid, which sediments at 75S and contains no RNA.(58–62) It is suggested that empty 75S particles serve as a storage form for particle formation, and can be dissociated *in vitro* into 14S pentamers to form new capsids.(63, 64) 14S pentamer subunits have RNA binding activity, but 75S particles do not, suggesting that the viral pentamers assemble around the RNA genome to form a provirion, and empty capsids are dead ends.(65–67) Following the incorporation of the RNA genome into the capsid structure, the provirion undergoes a maturation process, characterized by the internal cleavage of VP0 capsid protein, resulting in the VP2 and VP4 products.(68–70) This cleavage converts a provirion into an infectious viral particle. Both immature and mature virions sediment at 150S on a sucrose gradient, so it is not possible to distinguish the particles by buoyant density.(71)

For this reason, we employed radioactive labeling of newly formed EV-D68 particles, followed by sucrose gradient fractionation to separate radiolabeled empty and RNA-containing viral particles. We then visualized capsid proteins present in virion fractions by SDS-PAGE and autoradiography. Ammonium chloride, a non-specific acidification inhibitor that acts rapidly to block acidification of cellular compartments, was selected as the inhibitor for this work. Our previous publication on poliovirus maturation demonstrated that vesicle acidification promotes the maturation of the assembled poliovirus, converting the particles into infectious virions, so we employed a similar approach to EV-D68. Our data demonstrate that both EV-D68 capsid production and virus maturation occur independently of vesicle acidification (Figure 4A).(37) The size of 150S peaks of EV-D68 between 3-5.5 hpi was not altered by ammonium chloride treatment added at 0 hpi and continued throughout the infection (Figure 4A).

**Figure 4.**
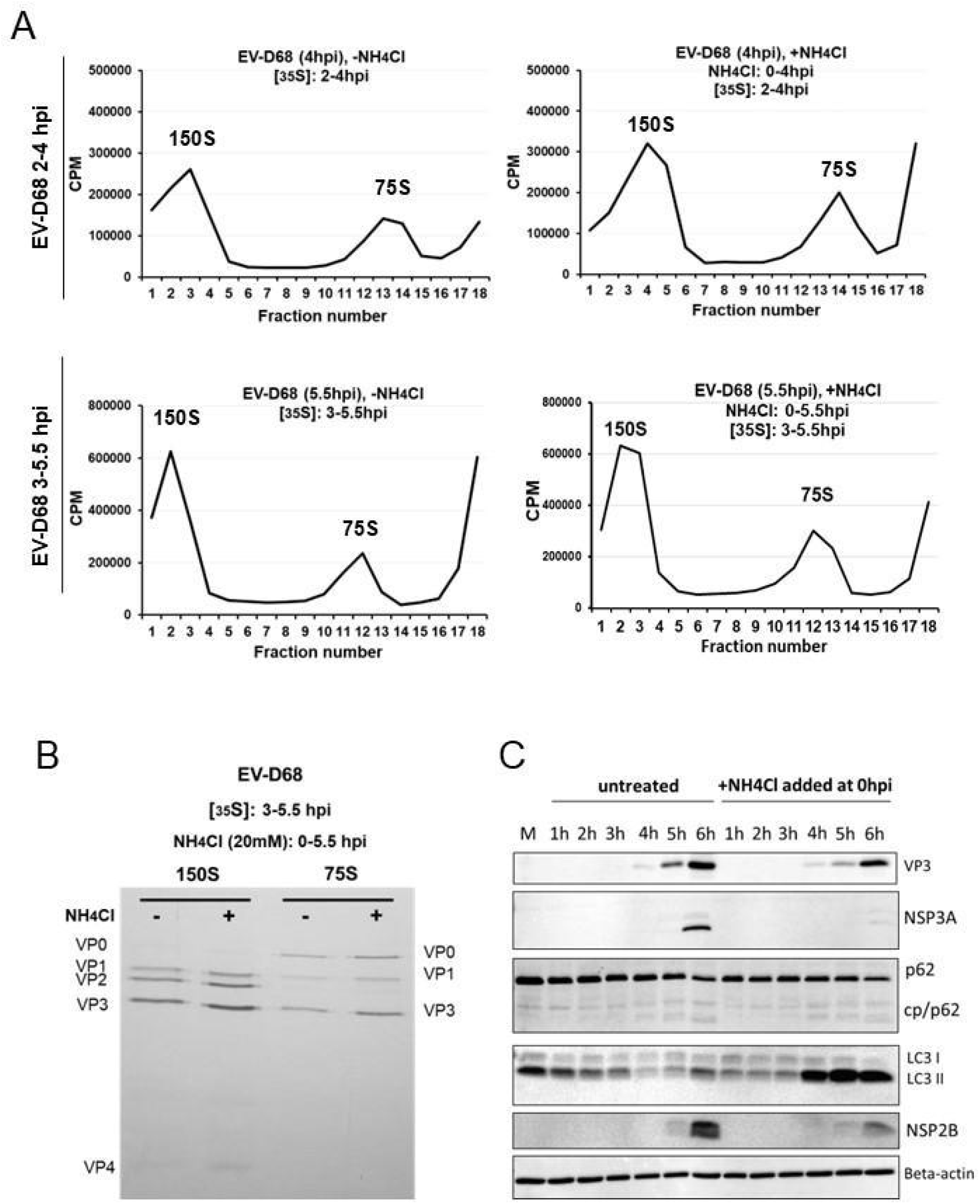
EV-D68 capsid production and virion maturation do not depend on cellular acidification, however, the production of several non-structural proteins does. (A) H1HeLa were infected at an MOI of 20, and the half of samples were treated with ammonium chloride added at 0hpi and continued until the end of infection (5.5 hpi). Cells were labeled with 35S-Methionine either from 2 hpi until collection at 4 hpi (A, upper panel), or from 3 hpi until collection at 5.5 hpi (A, lower panel). Cellular lysates were separated on 15-30% freshly prepared sucrose gradients and subjected to ultracentrifugation. Fractions were collected using the Fraction System and the counts per minute (CPM) were measured for each fraction. The experiments were independently repeated three times and the representative gradients are shown. (B) The collective three fractions of each determined peak (150S and 75S) were pooled and run on SDS-PAGE. The 35S-Methionine labeled bands were visualized using autoradiography films. The bands are labeled according to the expected relative migration pattern, while VP2 is identified by its absence in the 75S peak. (C) Production of several non-structural proteins of EV-D68 decreases upon de-acidification. H1HeLa cells were either untreated/mock (M) or infected with EV-D68 at an MOI of 20 for 6 h. At 0 hpi, the infected cells were washed, followed by either adding normal DMEM (EV-D68; -NH_4_Cl) or DMEM with 20mM of ammonium chloride (EV-D68; +NH_4_Cl), and samples were collected every hour during infection. Samples were subjected to western blot analysis for traditional autophagy markers: LC3B, p62(SQSTM1), and its Cp (cleavage product) and viral proteins: VP3 (virus structural capsid protein 3), as well as non-structural proteins, such as 2B (NSP2B) and 3A (NSP3A). Beta-actin served as a loading control.

Pooled 75S and 150S fractions from lysates collected at 5.5 hpi were analyzed by SDS-PAGE. We identified three labeled bands in 75S fractions in both treated and untreated cells, which we have labeled as VP0, VP1, and VP3 based on their expected relative mobility, while in the 150S fraction, we detected the VP2 band at its expected migration between VP1 and VP3 (Figure 4B). However, upon treatment with ammonium chloride, no changes in VP0/VP2 ratio are observed (Figure 4B), suggesting that, unlike PV, EV-D68 employs a strategy for virion maturation which is independent of low cellular pH. To confirm that these results were not specific to ammonium chloride treatment, we repeated the virus particle radiolabeling and sucrose gradient purification experiments using Bafilomycin A1 to inhibit vesicle acidification throughout the infection (Figure S3A). We observe no difference in VP0/VP2 ratio between Bafilomycin A1-treated EV-D68 fractions vs. untreated EV-D68 fractions (Figure S3B). The higher total viral-peak-associated radioactivity from Bafilomycin A-treated EV-D68 vs untreated might be explained by the above-mentioned delay in the infectious particle production. Interestingly, we noticed that the use of acidification inhibitors results in a distinct virus protein production pattern, as non-canonical stoichiometric ratios of several viral nonstructural proteins appear different upon our treatments in comparison to untreated infection (Figure 4C). To our surprise, only viral non-structural protein levels are affected by treatments, while ratios of the structural proteins remain unchanged (Figure 4C). We sought to investigate whether such a phenomenon occurs in the case of poliovirus infection, however, the same treatment led to no noticeable changes in PV protein levels, neither in structural, as evidenced by capsid VP3 expression, nor non-structural proteins 2C and 2B (Figure S3C), indicating that EV-D68 and PV rely on different strategies in their life cycle regarding cellular acidification.

### Both empty and RNA-containing virions are lost when vacuole acidification is inhibited

We wanted to separate acidification effects on viral entry from effects on the rest of the virus life cycle. Based on viral replication kinetics data and virus production growth (Figure 1), we define 2 to 4 hpi of the infection cycle as a logarithmic phase of EV-D68 infection, reflected in enhanced RNA replication and rapid production of functional viral particles. We hypothesize that midway through the logarithmic growth phase, when genomic RNA replication and packaging begins to decrease, compartments must acidify. vRNA replication depends on unique membrane replication organelles (ROs), the formation of which occurs between 2 to 3 hpi.(72) Thus, we wanted to know if the acidification of these organelles within the logarithmic phase is essential to EV-D68 production. We decided to test whether rapid de-acidification after the transition point affects virus yield.

Treatment of EV-D68-infected cells with ammonium chloride beginning at 3 hpi dramatically decreases viral titer (Figure 5A) and reduces genomic RNA levels (Figure 5B). Treatment with ammonium chloride beginning at 3 hpi has a critical effect on viral RNA levels, and the same treatment initiated post log phase (4 or 5 hpi) also significantly affects the levels of viral RNA. Finally, upon 3-6 hpi treatment with ammonium chloride, using radioactive pulse-labeling of viral proteins with ^35^S-Methionine followed by gradient sedimentation, we are unable to detect viral capsids (Figure 5C). While the absence of 150S RNA-containing peaks is not surprising since ammonium chloride inhibits vRNA levels, the absence of empty capsids at 75S is surprising. For EV-D68, even empty capsids do not form, indicating that vesicular acidification is necessary for forming empty capsids or for maintaining the stability of the capsids. In addition, we used confocal immunofluorescence microscopy (IFA) of EV-D68 infected H1HeLa cells to observe the effect of ammonium chloride treatment on viral infection. Infected cells were either treated with ammonium chloride (3 hpi to 6 hpi) or left untreated, and we monitored infection using dsRNA as a viral marker (Figure 5D). The number of infected cells at 3 hpi appears lower upon NH_4_Cl treatment than in an untreated infection. Moreover, upon treatment with ammonium chloride from 3-6hpi, viral dsRNA formed large agglomerate structures, localized in a perinuclear fashion, while no such structures are observed in untreated EV-D68-infected cells (Figure 5D, zoomed image).

**Figure 5.**
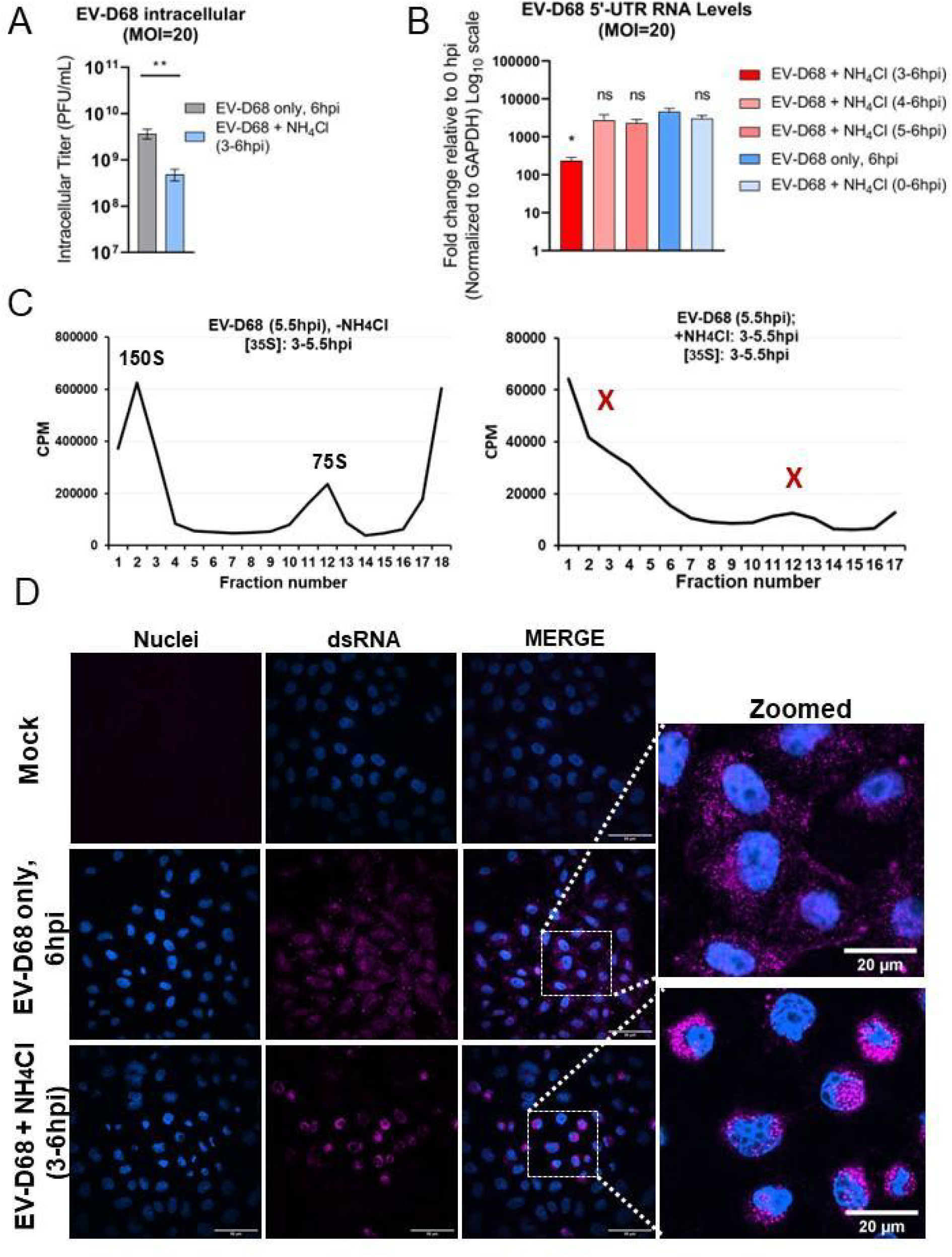
Vesicle acidification after the transition point of the EV-D68 cycle is essential for virion formation. (A) Titers of intracellular EV-D68 upon treatment with ammonium chloride (NH_4_Cl, 20 mM). H1HeLa were infected with EV-D68 at an MOI of 20; after 30 min of virus adsorption at 37°C (0 hpi), cells were washed twice to remove the initial virus input and the infection was allowed to run for 3h. At 3hpi ammonium chloride (20mM in complete DMEM) was added to the cells until the end of the infection (6hpi). Control samples (Infection only) were left untreated until virus collection at 6hpi. Intracellular titers were analyzed by plaque assay. Unpaired student’s t-test was used for the statistical analysis (***= p< 0.001; **= p< 0.01; *= p ≤ 0.05; ns=not significant). (B) Effect of ammonium chloride treatment on viral RNA levels assessed by qRT-PCR. Cells were infected with EV-D68 at an MOI of 20 for 30 min at 37°C, then the residual virus was washed away, followed by either adding normal media (untreated infection, EV-D68; -NH_4_Cl) or media containing 20mM of ammonium chloride (EV-D68; +NH_4_Cl) at different time points (at either 0 hpi, 3 hpi, 4 hpi or 5 hpi until virus collection (6hpi). Untreated infection (infection only, 6hpi) served as a control. Total RNA was extracted from all samples and subjected to qRT-PCR RNA analysis following cDNA synthesis. The level of GAPDH mRNA was used as an internal control. (C) Treatment with ammonium chloride within the transition point reveals the inability of the virus capsids to be formed. H1HeLa were infected at an MOI of 20, and the half of samples were left untreated (EV-D68 -NH_4_Cl) or treated with ammonium chloride (EV-D68 +NH_4_Cl) added at 3 hpi and continued until the end of infection (5.5 hpi). All cells were labeled with 35S-Methionine from 3 hpi until collection at 5.5 hpi. Cellular lysates were separated on 15-30% freshly prepared sucrose gradients and subjected to ultracentrifugation. Fractions were collected using the Fraction System and the counts per minute (CPM) were measured for each fraction. The experiments were independently repeated three times and the representative gradients are shown. (D) Confocal imaging of EV-D68 upon ammonium treatment during the transition point. H1HeLa were infected with EV-D68 (MOI=20) for 6h or left uninfected (mock). Infected cells were then left untreated (infection only) or treated with ammonium between 3 to 6hpi (EV-D68 + NH_4_Cl, 3-6hpi). Cells were fixed and then stained for dsRNA (purple) and nuclei (blue). The zoomed-in panels show the same subset of cells.

Since EV-D68 RNA levels appear to be altered by ammonium chloride treatment post 3hpi, we sought to investigate whether its antiviral effect would be comparable to that of the well-known potent viral RNA replication inhibitor guanidine chloride (GuaHCl).(73–77) Guanidine chloride blocks the initiation of negative-strand RNA synthesis by inhibiting the function of the viral 2C non-structural protein.(78–81) Our results show that ammonium chloride treatment of EV-D68 is comparable to guanidine chloride, as both effectively decrease intracellular EV-D68 titers (Figure S4A), viral RNA levels (Figure S4B) and protein synthesis (Figure S4C). However, we believe that as a nonspecific weak base, ammonium chloride may act more rapidly and broadly than guanidine chloride.

In addition, IFA on EV-D68-infected H1HeLa cells with or without ammonium chloride or guanidine chloride treatment demonstrated that guanidine chloride has not led to a formation of dsRNA agglomerates, unlike ammonium chloride (Figure S4C), highlighting that both treatments efficiently decrease vRNA levels, albeit by different mechanisms.

### Disrupted EV-D68 capsid assembly through myristoylation inhibition results in decreased RNA levels

Treatment with ammonium chloride from 3-6hpi causes both capsid and viral RNA levels to decrease. It is difficult to distinguish whether the loss of vRNA is due to lower RNA replication, or lack of capsids failing to protect the synthesized vRNA from cellular RNAses. We theorized that intact capsids protect already-produced vRNA until virion release, so to test this, we wanted to inhibit capsid assembly in a specific fashion. Nearly all picornaviruses require co-translational N-terminal myristoylation by cellular N-myristoyltransferases (NMTs) of the precursor capsid protein VP4 to ensure the successful capsid assembly, stability and, therefore, functional virus production.(82–88) Disrupting this process in poliovirus abrogates the viral assembly and results in little to no virus production.(89) We employed the pan-NMT inhibitor of picornaviruses assembly IMP-366 (DDD85646), a pyrazole sulfonamide originally developed against trypanosomal NMT, to assess its effect on EV-D68 capsid assembly, as well as whether the later might affect viral RNA load.(82, 90)

IMP-366 (5μM) treatment drastically reduced EV-D68 intracellular yield, and the effect of IMP-366 appears proportional to the duration of treatment (Figure 6A). Drug treatment alone did not have a significant effect on cell viability (Figure S5A). Next, we assessed the effect of an IMP-366 on viral RNA loads when co-treated with ammonium chloride (Figure 6B). The results suggest that disrupted viral assembly results in reduced EV-D68 RNA levels, indicating that, as in other picornaviruses, vRNA replication and virion assembly are tightly connected, and maintaining the stability of viral capsids is crucial for vRNA maintenance. Interestingly, when the treatments were applied together, viral RNA levels remained unchanged, indicating they may counteract one another in some way (Figure 6B).

**Figure 6.**
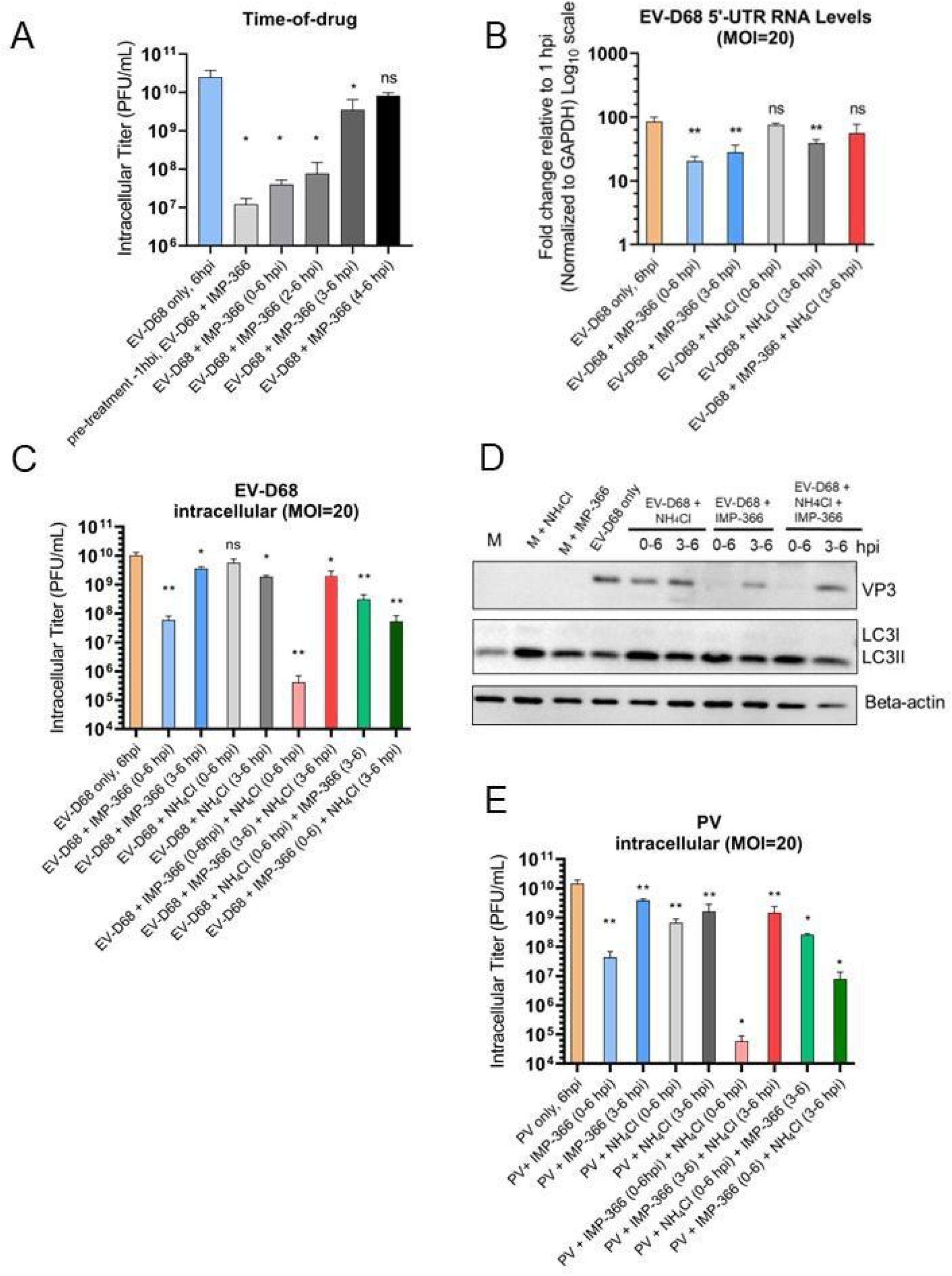
Disrupted EV-D68 capsid assembly through myristoylation inhibition results in decreased RNA levels. (A) Titers of intracellular EV-D68 upon treatment with ammonium chloride (NH_4_Cl, 20 mM), IMP-366 (5μM) or both at indicated time points via time-of-drug assay. H1HeLa were infected with EV-D68 at an MOI of 20; after 30 min of virus adsorption at 37°C (0 hpi), cells were washed twice to remove the initial virus input and the infection was allowed to run for 3h. At the indicated time points post-infection, ammonium chloride (20mM in complete DMEM) or IMP-366 (5μM) was added to the cells until the end of the infection (6hpi). Control samples (Infection only) were left untreated until virus collection at 6hpi. (B) Effect of ammonium chloride and IMP-366 on EV-D68 RNA levels assessed by qRT-PCR. Cells were infected with EV-D68 at an MOI of 20 for 30 min at 37°C, then the residual virus was washed away, followed by either adding normal media (EV-D68 only) or media containing 20mM of ammonium chloride (EV-D68; +NH_4_Cl), 5μM of IMP-366 (EV-D68; +IMP-366), or both at indicated time points (at 0 hpi or 3 hpi) until virus collection (6hpi). Untreated infection (infection only, 6hpi) served as a control. Total RNA was extracted from all samples and subjected to qRT-PCR RNA analysis following cDNA synthesis. Data represents relative expression of 5′UTR where samples are compared to the 1 hpi. All samples are normalized to GAPDH and have been log10 transformed. (C) Titers of intracellular EV-D68 upon treatment with ammonium chloride (NH_4_Cl, 20 mM), IMP-366 (5μM) or both at indicated time points. H1HeLa were infected with EV-D68 at an MOI of 20; after 30 min of virus adsorption at 37°C (0 hpi), cells were washed twice to remove the initial virus input and the infection was allowed to run for 3h. At indicated time point post-infection, ammonium chloride (20mM in complete DMEM) or IMP-366 (5μM in complete DMEM) was added to the cells until the end of the infection (6hpi). Control samples (EV-D68 only) were left untreated until virus collection at 6hpi. Intracellular titers were analyzed by plaque assay. (D) Effects of ammonium chloride and IMP-366 treatments added at different time points during EV-D68 infection on protein levels assessed by Western Blot. Cells were infected with EV-D68 at an MOI of 20 for 30 min at 37°C, then the residual virus was washed away, followed by either adding normal media (EV-D68 only), media containing 20mM of ammonium chloride (EV-D68+NH_4_Cl), media containing 5μM of IMP-366 (EV-D68 +IMP-366), or media with both treatments added simultaneously (EV-D68+NH_4_Cl+IMP-366), either at 0hpi or 3 hpi as indicated until final point of sample collection (6hpi). Untreated infection (EV-D68 only, 6hpi) served as a control. Samples were subjected to Western Blot analysis for autophagy marker LC3B and viral protein VP3 (virus structural capsid protein 3). Beta-actin served as a loading control. (E) Titers of intracellular PV upon treatment with ammonium chloride (NH_4_Cl, 20 mM), IMP-366 (5μM) or both at indicated time points. H1HeLa were infected with EV-D68 at an MOI of 20; after 30 min of virus adsorption at 37°C (0 hpi), cells were washed twice to remove the initial virus input and the infection was allowed to run for 3h. At indicated time point post-infection, ammonium chloride (20mM in complete DMEM) or IMP-366 (5μM in complete DMEM) was added to the cells until the end of the infection (6hpi). Control samples (EV-D68 only) were left untreated until virus collection at 6hpi. Unpaired student’s t-test was used for statistical analyses (***= p< 0.001; **= p< 0.01; *= p ≤ 0.05; ns=not significant).

To analyze whether the effect on RNA levels affects viral yield, we performed analysis of viral titers after treatments (Figure 6C). Simultaneous treatment with both ammonium chloride and IMP-366 beginning at the transition point again resulted in no effect, indicating an apparent neutralization effect of treatments when used in combination (Figure 6C). This effect was confirmed by Western Blot demonstrating restored levels of VP3 structural protein upon combined treatment (Figure 6D). We analyzed viral titers of the same treatments using PV infection, and the response to IMP-366 is similar to EV-D68. For PV, however, treatment with ammonium chloride 20mM alone (both 0-6hpi and 3-6hpi) significantly affected PV intracellular titers, as predicted. We attribute this effect to compartment acidification required for maturation of PV virions (Figure 6E). We compared the effect of IMP-366 continuous treatment (0-6hpi) throughout infection on EV-D68 and PV RNA levels (Figure S5B). We observed differences in PV and EV-D68 RNA levels at all time points; we interpret this to mean that vRNA production and/or stability, in the absence of capsid, is distinct between these two viruses.

### De-acidification after the transition point leads to structural changes in replication organelles (RO) and multi-vesicular bodies (MVB)

Based on the data in Figure 5D, we hypothesized that the effect of ammonium chloride on EV-D68 RNA levels may be due to changes or disruption of replication organelles (ROs). Using TEM microscopy, we observe both ROs and vesicles which appear to be of autophagic origin.(72, 91) Our TEM data identify RO complexes similar to those observed in PV during EV-D68 infection (Figure 7).(92) We also detect an increase of multi-vesicular bodies (MVBs) in infected cells, comprising separate compartments (“pockets”) of lighter electron density than that of the cytosol. These “pockets’’ appear to contain vesicles of various sizes. In untreated EVD68-infected cells, MVBs can be found both in proximity to the nucleus and the plasma membrane, however, in NH_4_Cl-treated (3-6hpi) infected cells, the MVBs localize more closely to the nucleus in a perinuclear area. With

**Figure 7.**
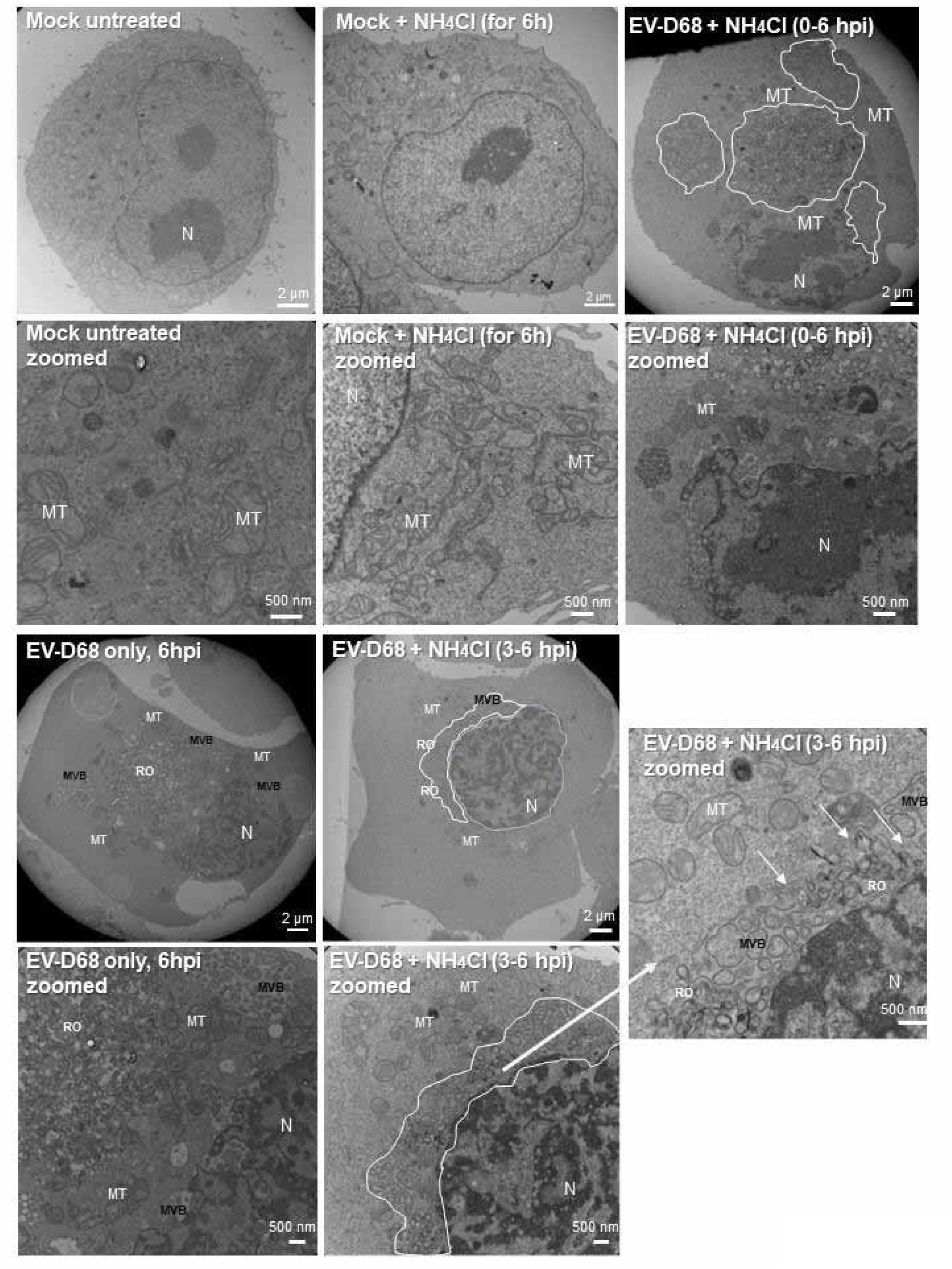
De-acidification during the “transition point” of viral infection leads to structural changes in replication organelles (ROs) and multi-vesicular bodies (MVB) Transmission electron micrographs of H1HeLa cells with mock infection or a 6h infection with EV-D68 (MOI = 20). H1HeLa were infected with EV-D68 at an MOI of 20; after 30 min of virus adsorption at 37°C (0 hpi), cells were washed twice to remove the initial virus input and the infection was allowed to run for 3h. At 3 hpi ammonium chloride (20mM) was added (EV-D68 + NH_4_Cl 3-6hpi) or infection left untreated (EV-D68 only, 6 hpi). Mock cells were either left untreated (mock) or treated with ammonium chloride for 6h (mock + NH_4_Cl for 6h). At 6h, all cells were fixed and subjected to TEM. Displayed images of the upper panel are followed by zoomed images of the same sample below, respectively. ROs: replication organelles; MVB: multi-vesicular bodies, MT: mitochondria; N: nucleus. Scale bar: 2μm or 500nm (zoomed images) as indicated. TEM images are representative of one biological experiment.

NH_4_Cl treatment post-transition point causes MVB compartments to appear disrupted, with their membranes no longer intact, and disrupted ROs and MVBs appear to merge (Figure 7, white arrows). The presence of ammonium chloride from the initiation of EV-D68 infection (added at 0 hpi, respectively) does not result in major disruption of MVB or RO compartments, highlighting the importance of this step in the transition from ROs to autophagosome-like vesicles for post-RNA virion formation.

## Discussion

The standard model of picornavirus replication is that the entire life cycle of the virus takes place in the cytoplasm. Here we show, in agreement with evidence from our group and others, that cellular compartments play a major role in virus production. Both PV and EV-D68 usurp the cellular autophagic machinery in order to generate functional virus progeny, and in both cases this process occurs independently from the canonical upstream signaling pathway.(41) However, the membrane-shaping machinery itself, including ATG7, which is required for lipidation of the LC3 protein and initiation of autophagosome formation, is required for efficient production of both viruses. This suggests a common feature of picornaviruses: using the membrane-shaping machinery of autophagy without engaging the nutrient- or stress-signaling machinery.

While the overall life cycle and relationship to autophagy are similar for PV and EV-D68, we were intrigued by the different responses of these viruses to cellular acidification inhibitors. Our lab has shown previously that blocking autophagosome formation inhibits PV RNA synthesis, while inhibiting vesicle acidification only affects maturation cleavage of PV particles.(37) In contrast, our current study demonstrates that prolonged treatment with several common inhibitors of acidification has no effect on generation of functional EV-D68 particles. To our surprise, the viruses demonstrated different kinetics: while treatment with acidification inhibitors abrogated the PV yields, EV-D68 responded to the treatment with an hour-long delay in virus production that recovered to normal yields by the end of a single-cycle infection (6hpi). For EV-D68, the *rate* of virus production is inhibited; for PV, the *extent* of virus production is reduced.(37)

This explained by our finding that, for EV-D68, acidification of cellular compartments is important at two distinct points of the viral cycle: cell entry, and after (but not *through*) the transition point from peak vRNA replication to virion maturation and egress. This is illustrated in our model (Figure 8). Acidification of the entry vesicle is a common mechanism for genome release. Data regarding pH requirements for PV entry have been inconclusive, but several studies have shown that acidification is important for EV-D68 uncoating.(57, 93–98) The delay in EV-D68 production is related to the first 30 min within viral entry into the cell, which suggests an effect on viral fusion with the entry vesicle.

**Figure 8.**
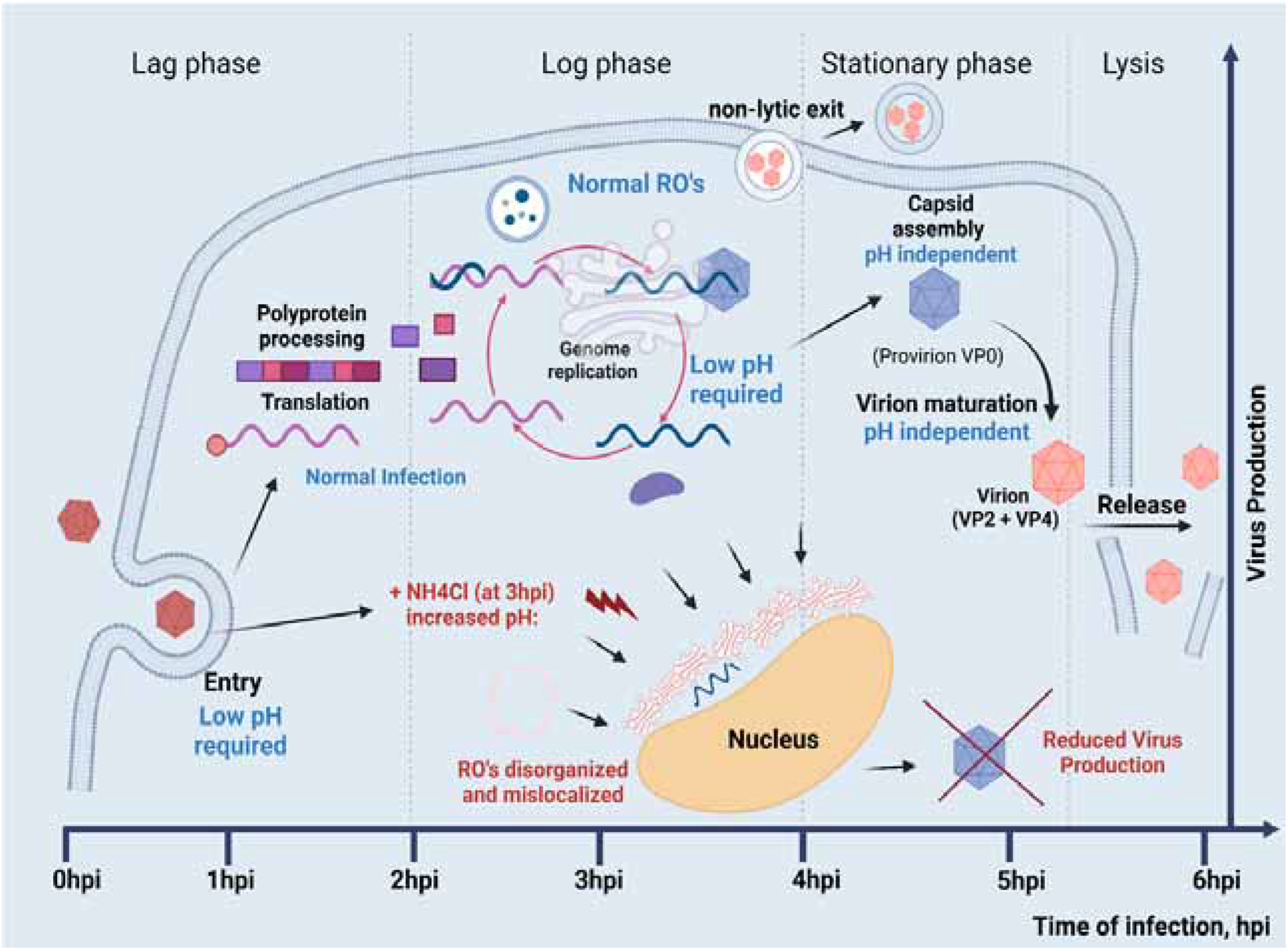
Model. Time of infection progresses left to right. Early in infection, low pH is required for EV-D68 entry. During the log phase of virus growth, low pH compartments are required for assembly and maintenance of both empty and genome-containing capsids. Low pH is also required for normal localization and morphology of viral replication organelles (ROs). From 3-4hpi, infections transition from the peak of vRNA replication, to EV-D68 capsid maturation and egress; after this transition point, low pH compartments are no longer required.

In our study both PV and EV-D68 required low pH for the early steps within viral entry into the host cell. The weak bases have this effect, although Brefeldin A with its specific activity against proton pumps did not. Low pH requirements for poliovirus entry could be explained by viral tolerance to low pH, as initial infection in the gut epithelia requires passing through the highly acidic environment of the stomach. However, it is surprising that EV-D68, as a virus primarily replicating in upper and lower respiratory epithelia, also required acidic pH early upon infection. On the other hand, several clinical studies reported that EV-D68 genetic material has been detected in wastewater and stool specimens, suggesting that potential fecal-oral transmission of the EV-D68 is also possible.(99–102)

Acidification after the transition point is critical for virion production and stability. We also find it causes changes in vRNA levels and in vesicular distribution in the cell. This differs from our previous findings for PV.(103) The mechanisms of picornavirus assembly and subsequent capsid maturation are not well understood. While poliovirus and coxsackie require myristoylation and cleavage of the VP0 capsid protein for efficient assembly, in several parechoviruses and kobuviruses VP0 is neither cleaved nor myristoylated.(82, 104) Here we show that for EV-D68, unlike PV, vesicle acidification does not play a crucial role in capsid maturation. Moreover, inhibition of myristoylation resulted in dramatic loss of intracellular viral titers, which suggests that, as for other picornaviruses, myristoylation is important for EV-D68 capsid stability.

Importantly, vesicle acidification is also required for EV-D68 genome maintenance throughout infection. EV-D68 vRNA levels are tightly linked to capsid formation and stability, in contrast to PV, whose RNA replication or capsid stability does not depend on low pH.(37) For EV-D68, it is possible that vRNA levels could be lowered by replication inhibition. However, we propose that lack of vRNA protection by capsids is to blame for loss of viral genomes late in infection. We hypothesize that viral replication organelles, which form in the cytoplasm during the early log phase of viral infection, require membrane potentials for their proper generation, structural integrity, and, importantly, localization. De-acidification of cellular compartments via rapid deprotonation with ammonium chloride leads to morphological changes in the RO, suggesting that intact organelles are crucial for virus production. More importantly, we show that timing within which the interruption of such organelles occur is critical. The mid-log phase transition point is a vital step, when the majority of virus activity switches from viral RNA replication to capsid maturation and egress. Our data suggest that rapid inhibition of acidification during this phase leads to an inability to form stable capsids, and a decrease of viral RNA levels.

For the first time, we demonstrate that acidification of cellular compartments within specific phases of the life cycle of a picornavirus, defined as before and after the transition point, has different outcomes during infection (Figure 8). Continuous treatment throughout infection with acidification inhibitors results in reduced levels of several non-structural proteins upon EV-D68 infection. No similar decrease in non-structural protein levels has been observed from poliovirus. Since picornavirus proteins are produced from a single polyprotein and are initially at equal stoichiometry, we propose that loss of individual non-structural proteins is due to instability and degradation in the absence of normal replication organelles.

Picornavirus replication is often described as being entirely cytosolic.(105, 106) However, multiple studies have now demonstrated the importance of membrane potentials on various stages of the picornavirus life cycle. Here we show that inhibiting acidification disrupts normal formation and localization of EV-D68 replication organelles. Moreover, our study contributes to understanding differences in the life cycles of two picornaviruses, EVD68 and PV, whose distinct initial tropisms belie similar neurological outcomes.

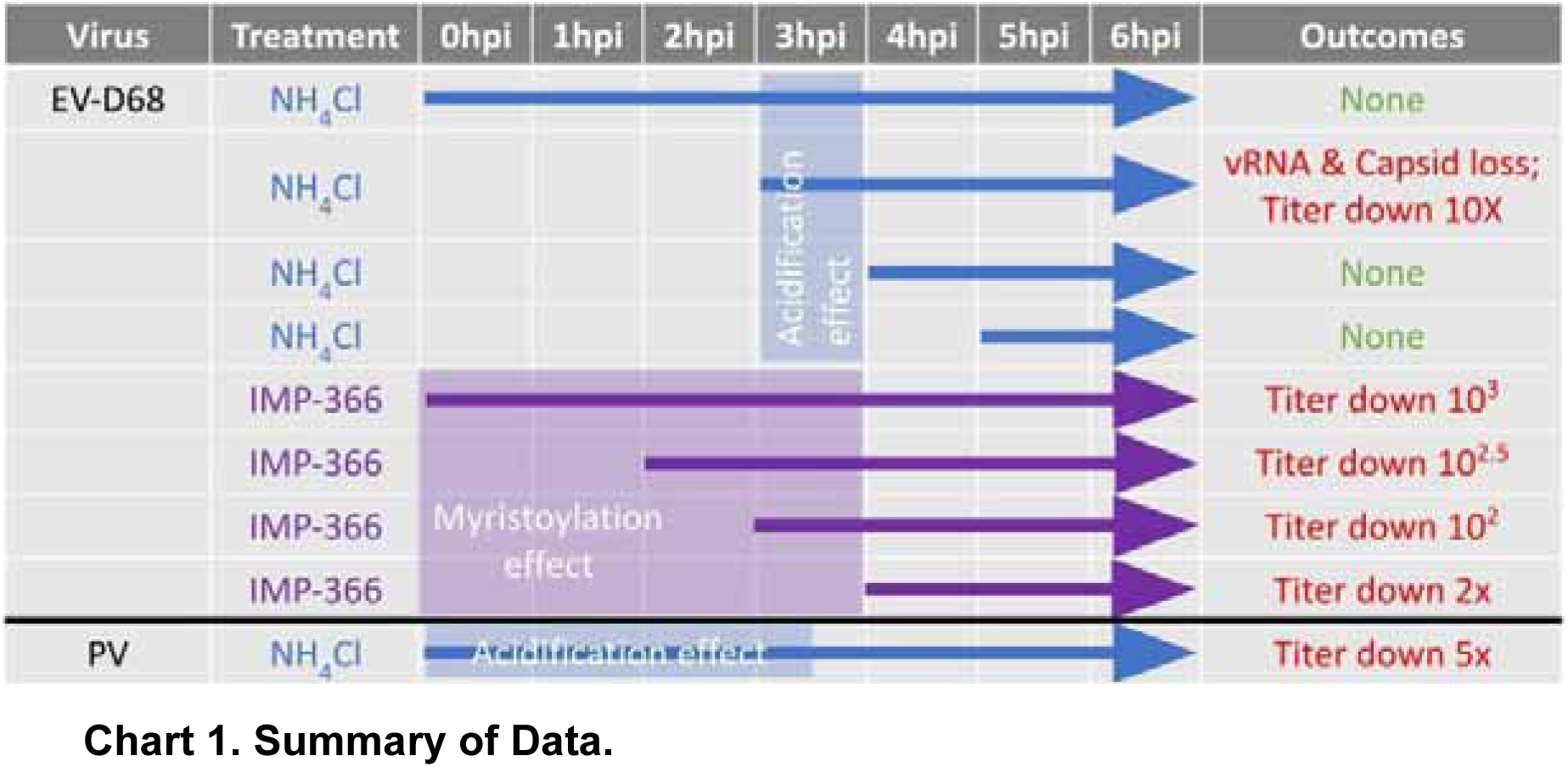

## Materials and Methods

### Cells

H1HeLa (Human cervical adenocarcinoma cells) were maintained in Dulbecco′s Modified Eagle′s Medium (DMEM; Gibco) supplemented with heat-inactivated 10% fetal calf serum (FCS; Gibco), sodium pyruvate (1X; Gibco), and penicillin/streptomycin solution (1X; Gibco). Cells were grown at 37°C, 5% CO_2_.

### Viruses and plasmids

To avoid accumulating viral mutations upon passaging, EV-D68 (prototype Fermon) or Poliovirus (type 1 Mahoney) were directly prepared from corresponding plasmids EV-D68 Fermon or pGEM encoding full-length cDNA copies of viral genomes as described below.

### Bacterial transformation

50ng of each plasmid containing viral cDNA genome were used for transformation of competent bacteria (*E. coli* strain DH5α, Thermo Scientific) by heat-shock, immediately followed by recovery 45-min incubation in Super Optimal broth with Catabolite repression (SOC) medium at 37°C with shaking (220 rpm). The transformed cells were spined at 3000g for 5min, the supernatant was discarded, and the bacteria pellet was gently resuspended in SOC media. Then the transformed bacterial suspension was plated on agar plates containing ampicillin (50mg/ml) and incubated overnight at 37 °C. The next day individual colonies were picked and transferred for expansion growth in 5mL or 200mL of LB broth with ampicillin (50mg/mL) in a shaker for overnight incubation at 37 °C. The overnight culture was used either for freezing glycerol stocks (40%) in cryovials for storage at −80 °C until use or for maxi-prep plasmid DNA extractions.

### Plasmid isolation and quantification

Isolation of EV-D68 Fermon and pGEM plasmids from transformed bacteria were performed using the EndoFree QIAGEN Plasmid Maxi Kit (12362; Qiagen) according to the manufacturer’s instructions. Final DNA concentration measurements were performed using the DeNovix DS-11 spectrophotometer (SN-06428, USA).

### Plasmid linearization

Viral genome-containing plasmids were then linearized via restriction digestion. The digest reaction total volume of 20µl with 1 or 2µg of the desired plasmid in a suitable buffer was set up. Restriction *EcoRI* enzyme (ER0271, Thermo Scientific) was used for pGEM plasmid, while *MluI* restriction enzyme (ER0561, Thermo Scientific) was used for EV-D68 Fermon in the equivalent of 1U of restriction enzyme/1µg of plasmid DNA. The reactions were let incubated overnight at 37°C. The linearized DNA was precipitated by adding 1µl of 0.5M EDTA (pH 8.0), 2µl of 3M sodium acetate (pH 5.2), and 50µl of EtOH (100%) followed by incubation at -20°C for over 15 minutes and a 20-min spin at 15,000 rpm, 4°C. The supernatant was carefully aspirated, and the dry DNA pellet was resuspended in nuclease-free water. 1:10 dilution of the reaction product was used to measure DNA concentration. Plasmid digestion was confirmed via the separation of linear DNA fragments via agarose gel electrophoresis (0.8∼1.2%(w/v)) in TAE buffer (1x; 40mM Tris, 20mM acetic acid, 1mM EDTA). Linearized templates were used for the following *in vitro* RNA transcription.

### In *vitro* RNA transcription and virus stock preparation

A total of 1μg either of pGEM plasmid containing a cDNA copy of the entire poliovirus type 1 Mahoney genome or 1ug of EV-D68 Fermon-encoding entire genome plasmid was transcribed using the Megascript T7 transcription kit (AM1334; Thermo Scientific) according to the manufacturer’s instructions. Total RNA concentration measurements were performed using the DeNovix DS-11 spectrophotometer (SN-06428, USA) and stored at -70°C until use. 12ug of the RNA product in serial dilutions (10µg, 1µg, 100ng, 10ng, 1ng, and 0.1ng RNA) was transfected into H1HeLa cells using 24µl of Lipofectamine 2000 (11668019, Thermo Scientific) transfection reagent and OptiMEM medium (Gibco) in a total volume of 600µl per reaction. 500µl of the reaction mixture was added to the cells, incubated for 4h at 37°C, and then overlaid with 2% bactoagar (214010, Becton, USA) in MEM (2X, Gibco). Plaque formation was allowed for 36-48 hours, then individual plaques were picked using a glass pipette and resuspended in 250µl of phosphate-buffered saline (PBS). After 3× freeze/thaw cycles the virus was released from the plaques and added to 1 mln of H1HeLa cells seeded previously in a 60 mm cell culture dish. At 6 hours post-infection (hpi) the media was removed, and cells were scraped into 500µL of PBS, followed by lysis via similar 3× freeze/thaw cycles. The lysate was then added to 6×10^6^ H1HeLa cells in a 10 cm cell culture dish, and at 6hpi the post-infection media was removed, and cells were collected in 1mL PBS. Cells were lysed by 3× cycles of freeze/thawing, cell debris was removed by centrifugation at 6000×g for 3 minutes at room temperature. The supernatant was transferred to a fresh 1.5mL Eppendorf tube and the titer of the virus was determined via titration on H1HeLa cells as described below.

### Virus infection and titration

Infections were performed in PBS with an adsorption time of 30 min at 37°C unless specified otherwise. The appropriate amount of virus was calculated as MOI in accordance with seeded cell density and concentration of the viral stocks (PFU/mL). The thoughtful removal of virus inoculum, followed by 2xPBS washes and the addition of DMEM complete medium after adsorption was considered as 0 hpi time point of infection. Infections were performed at MOI of 0.1 (low) or 20 (high) and were carried out for 6h unless specified (5.5 hpi for radioactive labeling experiments) and collected in 1 mL of PBS. All virus stocks were stored at -80°C at all times.

Titrations of the intracellular virus were carried out via 3× freeze/thaw lysis of the infected cells in PBS solution, followed by a short spin of 3000×g for 5 minutes at room temperature. The lysate was added in serial dilutions to a monolayer of H1HeLa cells (approx. 90% cell confluency) for a 30-min absorption at 37°C, after which cells were overlaid with bactoagar (2%) in MEM (2X; Gibco). Each dilution was plated in triplicate. Plaques were allowed to develop for 36-48 hours, then the bactoagar overlay was removed, cells stained with crystal violet (1%) for 3 min, and plaques counted. Dilutions were adjusted for dilution factor and reported as PFU/mL. Titrations of extracellular viruses required a short 3000xg 5-min spin to remove the cells from the supernatant and followed a similar infection protocol.

### Drug treatments

Ammonium chloride (NH_4_Cl, Sigma-Aldrich) at 20mM, Bafilomycin A1 (Baf A1; Cell Signaling) at 0.1µM, and chloroquine (CQ; Sigma-Aldrich) at 100µM were added to complete DMEM or DMEM without Methionine (Gibco). The media were then added to cells after viral adsorption (0 hpi) or at different time points of infection as specified. Guanidine hydrochloride (GuaHCl, G3272, Sigma-Aldrich) was used as an inhibitor of viral replication at 2mM.

DDD85646/IMP-366(2,6-dichloro-4-[2-(1-piperazinyl)-4-pyridinyl]-N-(1,3,5-trimethyl-1H-pyrazol-4-yl)-benzenesulfonamide) was purchased from Cayman Chemicals and used as an inhibitor of viral assembly at the final concentration of 5 µM. The cells were subsequently collected for Western blot, RNA isolations, plaque assays, or fixed with PFA (4%) for confocal imaging.

### RNA isolation, cDNA synthesis, and qRT-PCR

Total RNA was isolated from H1HeLa cells using TRIzol (15596026, Ambion) according to the manufacturer’s instructions. Thermo Scientific RevertAid H Minus First Strand cDNA Synthesis Kit (K1632, Thermo Scientific) was used to synthesize the cDNA following the removal of the genomic DNA using the DNAse I (EN0521, Thermo Scientific). qPCR was performed using KiCqStart SYBR qPCR Ready Mix (KCQS1, Sigma-Aldrich) with the 7500 Fast Dx Real-Time PCR Instrument (Applied Biosystems) according to the manufacturer’s protocols. The following primers, 5′ TAACCCGTGTGTAGCTTGG-3′ and 5′-ATTAGCCGCATTCAGGGGC-3′, which are specific to the 5′ untranslated region (UTR), were used to amplify EV-D68. GADPH primers were used for the control. The gene expression results were normalized to GAPDH and plotted as relative expression compared to the 1h infection-only time point.

### Virus entry assays

The synchronized infection assay on early events of viral attachment and entry/fusion, as well as drug treatments’ impact on cell-free virus particles (virucidal neutralization assay), was carried out according to protocol described(56). A low MOI infection of 0.1 was used for all entry assays and all cells were collected at 6hpi and subjected to plaque assays as described above.

In the virucidal neutralization assay, the viral particles were incubated with a specified treatment at 37°C for 1h, then the mixture was diluted 20-fold to a subtherapeutic concentration of the drug, and the virus inoculum was subsequently added to H1HeLa cells. The 20-fold dilution served to titrate the treatment drugs below their effective doses and prevent meaningful interactions with the host cell surface. After adsorption for 60 min at 37°C, the unbound virus was washed twice with PBS (1x). The DMEM was added to the cells until the end of the infection carried on at 37°C for 6h.

In the virus attachment assay, the virus was added to cells at an MOI of 0.1, and adsorption was carried out for 1h at 4°C. After adsorption, cells were washed twice with ice-cold PBS (1x) to wash off the unbound virus and DMEM was added to the cells until the end of infection further carried out at 37°C.

In the virus entry/fusion assay, the adsorption of the virus was allowed for 1h at 4°C, then the unbound virus was washed twice with ice-cold PBS and the treatment at an indicated concentration was added to cells in DMEM only for 30min incubation at 37°C. After 30min the cells were additionally washed with PBS and DMEM was added until the end of infection at 37°C.

### CRISPR/CAS Knockouts

H1HeLa CRISPR cell lines exhibiting ATG7, RB1CC1, and PI3K3C KO’s were generated in the Translational Laboratory Shared Services CRISPR Core (TLSS-CRISPR) using the CRISPR-Cas9 mechanism with two synthetic single-guide RNAs (sgRNAs) per gene. CRISPR-Cas9 KOs were produced by nucleofection on the Lonza Amaxa™ 4D-Nucleofector platform. KO’s were confirmed by subjecting cells to PCR and Sanger sequencing, and subsequent sequence analysis using the Synthego ICE analysis platform.

### SDS-PAGE and Western Blot

Cells were collected in PBS, centrifuged at 6000 × g for 5 min, and resuspended in Radioimmunoprecipitation assay buffer (RIPA) buffer (MFCD02100484, Sigma-Aldrich), supplemented with Roche cOmplete Tablets Mini Protease Inhibitor Cocktail (11836153001, Sigma-Aldrich). Lysis of cells was performed on ice for at least 1h. Insoluble debris was spined at 6000x g for 5 min, cell extracts were subjected to Bradford assay for protein concentration measurements. Equal input of the protein was loaded onto 15% sodium dodecyl sulfate-polyacrylamide (SDS-PAGE) gel, run at 80V-110V, and then transferred for 1h at 100V to polyvinylidene fluoride (PVDF) membranes (ThermoScientific). The membranes were blocked for 1h with 5% skim milk-TBST and incubated with primary antibodies at 1:1000-1:2000 overnight on a shaker at 4°C. The membranes were re-washed 3× with TBST(1X) and incubated with the appropriate secondary antibody (1: 2000) (Goat Anti-Mouse IgG (H+L)-HRP Conjugate, Bio-Rad #1706516 or Goat Anti-Rabbit IgG (H+L)-HRP Conjugate, Bio-Rad #1706515) at room temperature for 1 h on a shaker before being developed by Western Lightning Plus-ECL (Bio-Rad) at 1:1 ratio. Images were acquired using ChemiDoc (Bio-Rad) software and Bio-Rad Image Lab (version 6.1) program.

### Radioactive protein labeling

H1HeLa (6 mln cells/10 cm culture dish) were infected at MOI of 20 PFU/cell. At the defined time indicated in each experiment, complete DMEM was removed from cells, cells were washed twice with methionine-free DMEM and labeling media was prepared by adding 75 uCi of [35S] methionine per mL (PerkinElmer, NEG709A) of methionine-free medium (Gibco). At the time of collection, cells were washed twice with methionine-free DMEM, scraped into 1ml of PBS, and centrifuged for 6000 × g for 5min. Radiolabeled cells then were resuspended and lysed using the previously described method. Proteins were separated in 15% SDS-PAGE gel. The gel was dried for 2h at 60°C using Gel Dryer 583 (Bio-Rad) and visualized on developing film overnight at room temperature.

### Radioactive labeling of viral particles

The labeling method was modified from the original protocol described(107). H1HeLa at 6 mln/10cm culture dish were infected at MOI of 20 PFU/cell. Labeling media was prepared by adding 100μCi [_35_S] methionine per milliliter of methionine-free DMEM (Gibco). At specific time points indicated in each experiment, the cells were washed 2x with methionine-free DMEM, and the labeling media was added to each dish. Depending on the time of interest, newly synthesized [_35_S] methionine-containing proteins were labeled. After labeling, cells were washed with PBS and scraped into 1mL of PBS. Cell suspensions underwent 3× freeze/thaw cycles, releasing the viral particles, followed by a short 6000g spin to get rid of the cell debris. The lysates were stored at - 80℃ until loaded onto gradients.

### Separation of radioactive-labeled particles via density centrifugation and fractionation

The method of separation of viral particles was modified from the original protocol described (107). Sucrose gradients were prepared fresh before each fractionation with 15% (w/v) and 30 (w/v)% sucrose in 10 mM Tris (pH 7.4)-10 mM NaCl-1.5 mM MgCl2. The Gradient Master gradient maker (Biocomp) was used for continuous 15-30 % gradient preparation. Gradients were prepared in Beckman Ultra-Clear Centrifuge Tubes (14 × 89 mm, Beckman Coulter). Following preparation, 1 mL of radiolabeled lysate was loaded on top of the continuous sucrose gradient and was sedimented via centrifugation for 3h at 27500 rpm at 15°C using a Beckman SW41 rotor in Beckman ultracentrifuge at slow acceleration and deceleration with no brake. After centrifugation, each tube was placed into the Fraction Recovery System (Beckman Coulter) and the bottom of the tube was punctured, 500µL fractions were collected from the bottom of the tube into 1.5mL Eppendorf tubes by gravity flow. Radioactivity in each fraction was calculated by taking 20µL of each fraction and diluting in 4mL of CytoScint Liquid Scintillation Cocktail (MP Biomedicals) with the following measurement in a liquid scintillation counter (Beckman LS-3801).

### Separation of radioactive-labeled viral proteins

Pools (x3) from peak fractions were collected, boiled for 5 min at 95°C, and loaded on 15% SDS-PAGE gel. The gel was fixed overnight at 50% methanol-10% acetic acid solution at 4C while shaking. After that the gel was washed with H_2_0 and soaked in radioactive enhancer solution (6NE9741, PerkinElmer) for 15min, followed by a wash with water. Then the gel was placed on filter paper, covered with plastic wrap, and subjected to drying for 2h at 60°C. The gel was placed on the developing film and exposed for 24-48 h at RT, and images of distinct radioactive bands were obtained using a Bio-Rad developing machine.

### Confocal microscopy

Cells were seeded on 18mm coverslips a day before infection. After the infection, coverslips were fixed in 4% paraformaldehyde for 20min at RT followed by three PBS(1X) washes. The samples were then blocked with a blocking buffer (3% FCS/PBS (1X)) for 1h at RT while on a shaker. Then the primary antibodies were added to each sample and incubated overnight at 4C. The primary antibodies used included: anti-VP3 (MA5-18206, Thermo Fisher), anti-dsRNA (MABE1134, Sigma-Aldrich) at 1:200 dilution. The following day, the coverslips were washed three times with PBS(1X) for 5min each and the secondary antibodies (Alexa Fluor 488 or Alexa Fluor 594) were added for 1h at RT. Before mounting on microscope slides, cells were additionally washed with PBS(1X). ProLong Glass Antifade Mountant (P36985, Thermo Fisher) with NucBlue was used and the coverslips were allowed to dry overnight before storing at 4C. Images were acquired either using the Echo Revolve microscope or W2 Olympus confocal microscope and analyzed with Fiji software.

### Transmission Electron Microscopy (TEM)

Samples were fixed in 2.5% glutaraldehyde, 3 mM MgCl_2_, in 0.1 M sodium cacodylate buffer, pH 7.2 overnight at room temperature. After buffer rinse, samples were postfixed in 1.5% potassium ferrocyanide, 2% osmium tetroxide in 0.1 M sodium cacodylate (2 hr) on ice in the dark. Following a dH2O rinse, samples were en bloc stained with 2% Uranyl acetate (UA) before dehydration through a graded series of ethanol. Samples were embedded in EPON and polymerized at 60°C overnight. Thin sections, 60 to 90 nm, were cut with a diamond knife on a Leica Ultracut UCT ultramicrotome and picked up with 2×1 mm formvar coated copper slot grid. Grids were observed on a Hitachi 7600 TEM at 80kV and images captured with an AMT CCD XR80 (8 megapixel camera - side mount AMT XR80 – high-resolution high-speed camera).

### Statistical analysis

The statistical analysis was performed using GraphPad Prism software (Version 7.03). Unless otherwise indicated, values represent the mean ± standard error of the mean (SEM) of at least 3 independent experiments. Samples were analyzed using a Student t-test for comparison, and statistical significance was set at a p-value of < 0.05.

## Acknowledgments

We are grateful to the Jackson, Frieman, and Coughlan lab members for discussions on the project. We thank Brandon Cooper, Andrea Casildo, and the Translational Core facility at the University of Maryland Baltimore School of Medicine for help in the generation of the CRISPR/CAS KO cell lines used in the study. We also thank the Confocal Microscopy Facility of the University of Maryland Baltimore School of Medicine for providing confocal imaging equipment and software. We additionally thank Dr. Barbara Smith and the Microscope Facility staff at the Johns Hopkins University School of Medicine for preparing TEM samples and acquiring images in this study. This work was supported by NIH/NIAID grants R01AI141359 and R01AI104928 to W.T.J.

---

**Figure S1.**
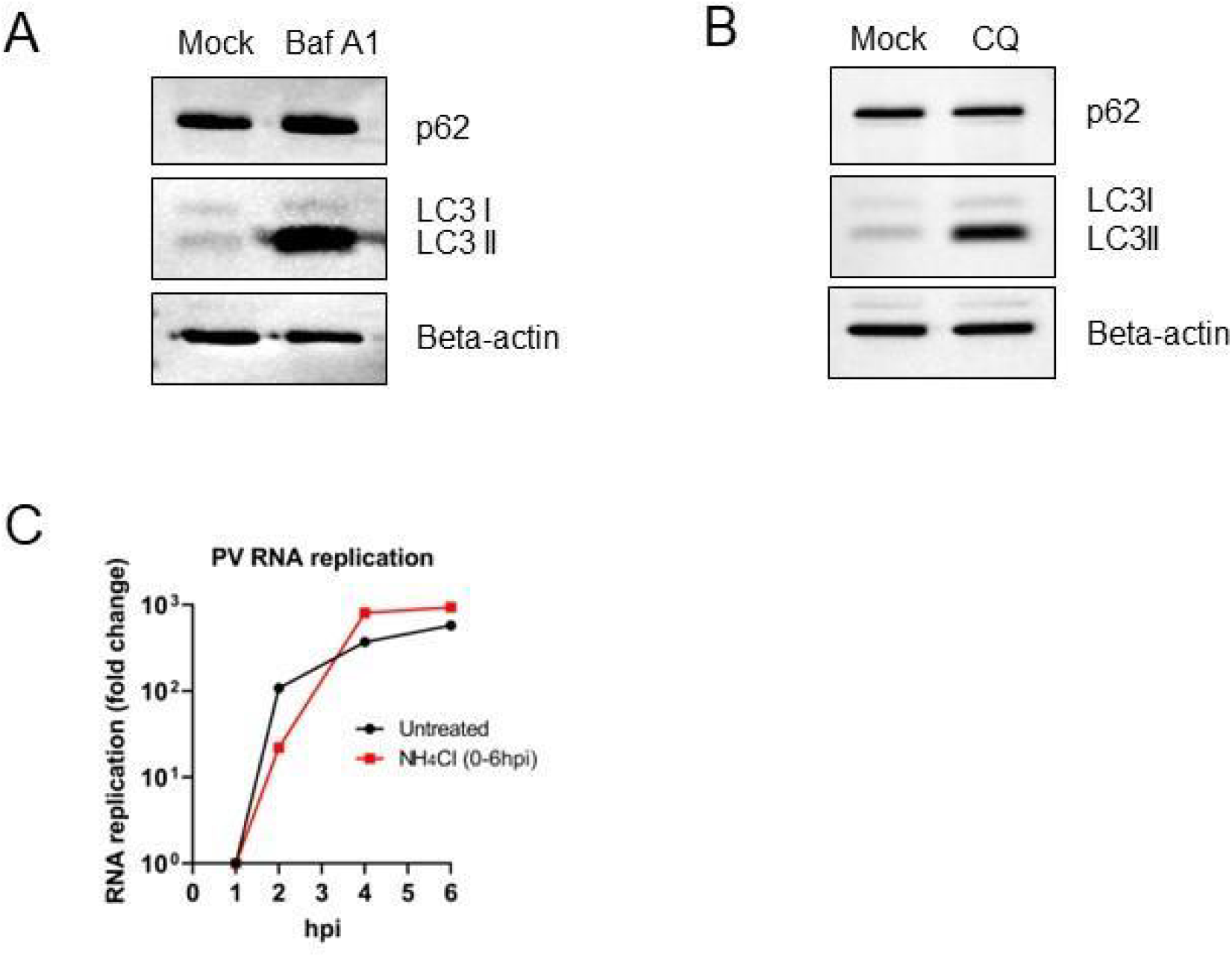
Related to Figure 2. Effect of the acidification inhibitors on autophagy markers. Autophagy markers SQSTM1 and LC3B in mock samples of H1HeLas, untreated or treated with chloroquine (CQ; 100uM) (A) or Bafilomycin (B) for 6h, were analyzed by Western blot. Beta-actin served as a loading control. The gradual accumulation of LC3B served as an indicator of blocked autophagy. (C) Effect of ammonium chloride treatment on PV RNA synthesis assessed by qRT-PCR. Cells were infected with PV at an MOI of 20 for 30 min at 37°C, then the residual virus was washed away, followed by either adding normal media (untreated infection, PV) or media containing 20mM of ammonium chloride (PV+NH_4_Cl). Cells were then collected at the indicated time points after infection, and total RNA was extracted and subjected to qRT-PCR RNA analysis following cDNA synthesis. The level of GAPDH mRNA was used as an internal control.

**Figure S2.**
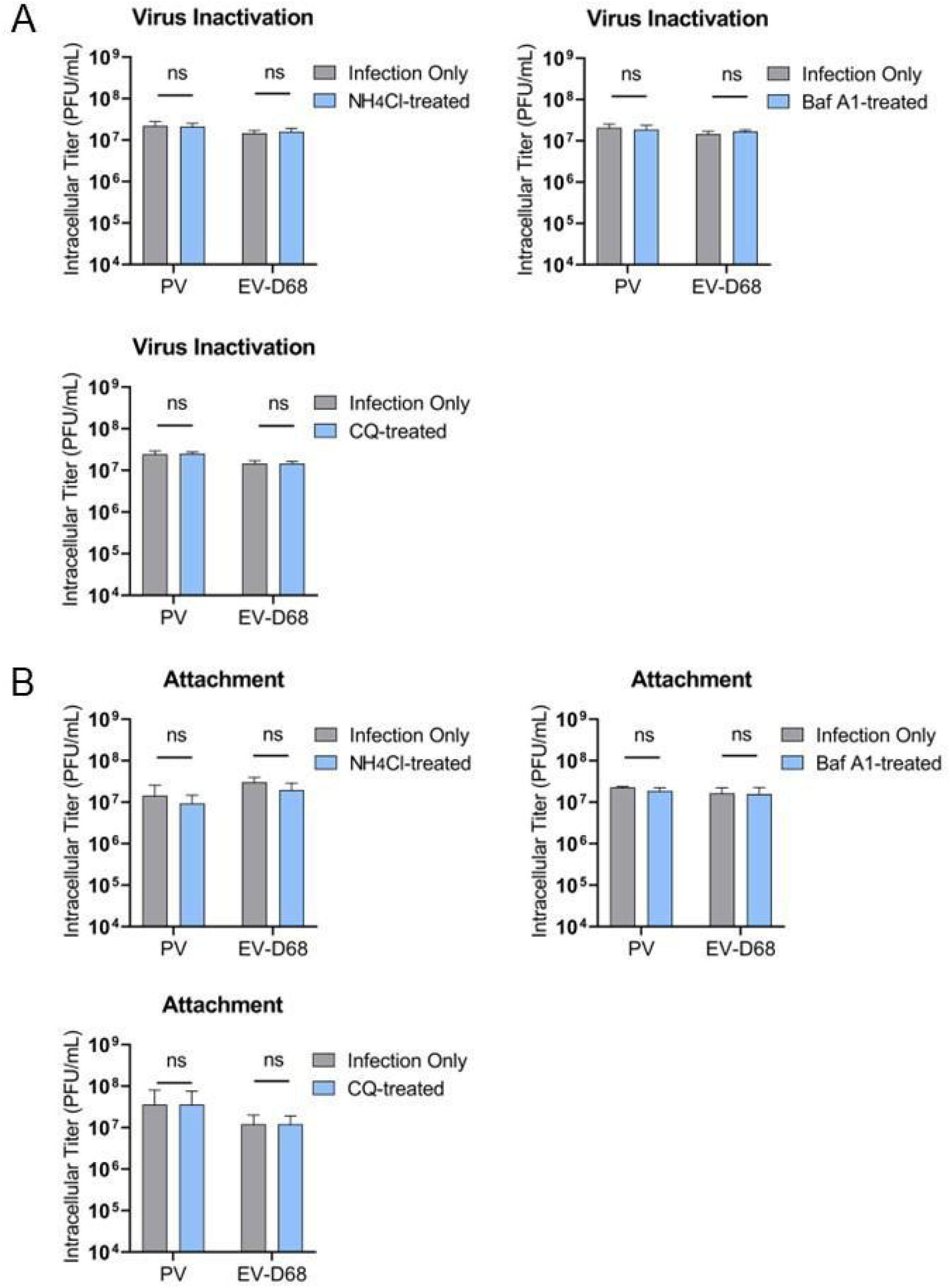
Related to Figure 3. Neutralization and attachment assays using acidification inhibitors. (A) Effect of acidification inhibitors on virus particle neutralization. Viruses and each drug were incubated for 1h at 37°C. Dilution of 20x of each drug was performed to subtherapeutic concentration to prevent drug-related alteration of cellular functions. (B) Effect of acidification inhibitors on viral attachment. H1HeLa were pre-chilled before the infection of PV or EV-D68 at MOI 0.1. The virus binding/attachment step was allowed to run for 1h at 4°C. Then the residual virus was washed at least twice and the cells were moved at 37°C for 6 hpi.

**Figure S3.**
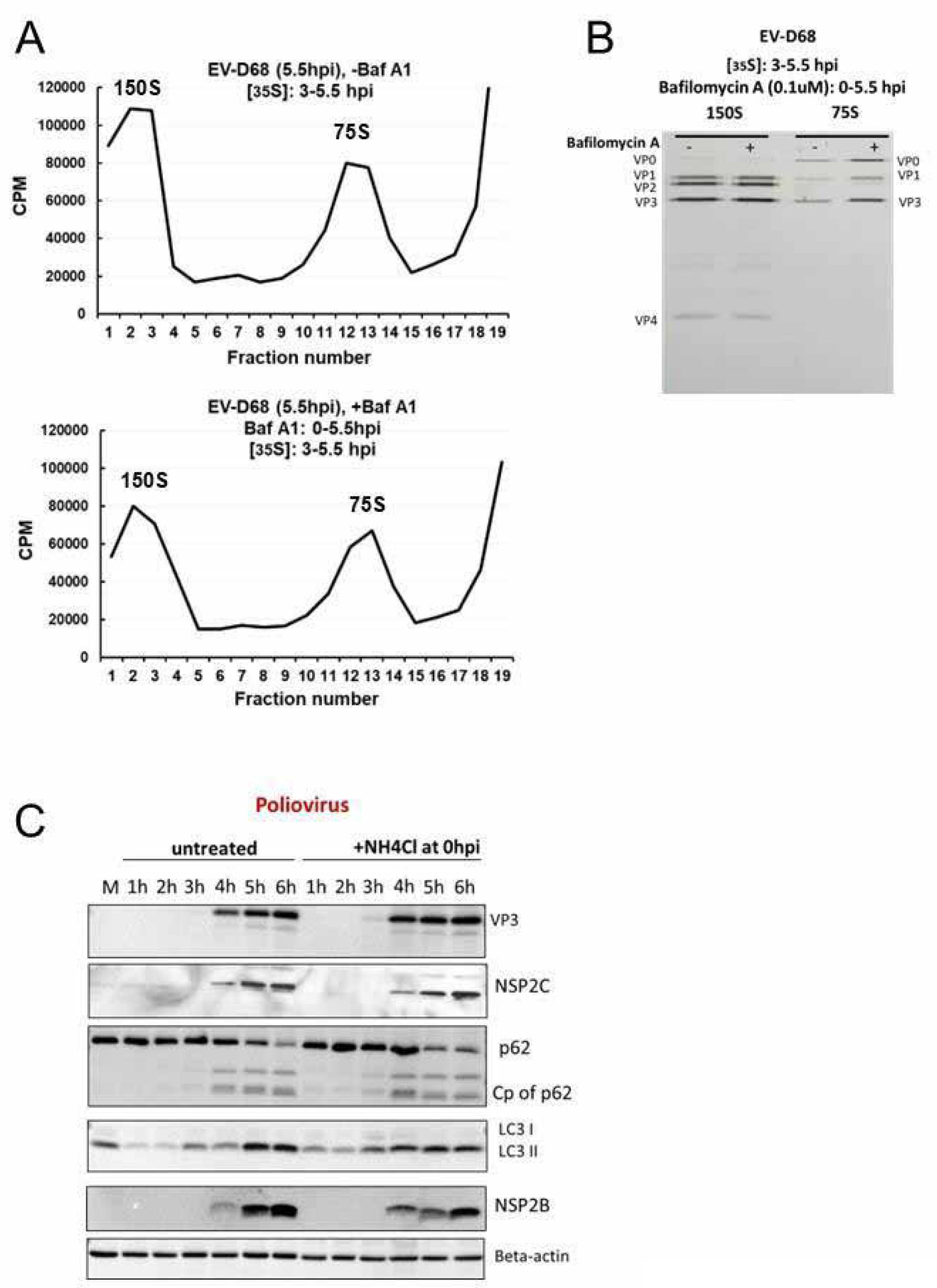
Related to Figure 4. EV-D68 capsid production and virion maturation is not altered by treatment with Bafilomycin A. (A) H1HeLa were infected at an MOI of 20, and the half of samples were treated with Bafilomycin A (0.1uM) added at 0 hpi and continued until the end of infection (5.5 hpi). Cells were labeled with 35S-Methionine either from 3 hpi until collection at 5.5 hpi (A). Cellular lysates were separated on 15-30% freshly prepared sucrose gradients and subjected to ultracentrifugation. Fractions were collected using the Fraction System and the counts per minute (CPM) were measured for each fraction. The experiments were independently repeated three times and the representative gradients are shown. (B) The collective three fractions of each determined peak (150S and 75S) were pooled and run on SDS-PAGE. The 35S-Methionine labeled bands were visualized using autoradiography films. The bands are labeled according to the expected relative migration pattern, while VP2 is identified by its absence in the 75S peak. (C) Production of several non-structural proteins of PV is not affected by treatment with acidification inhibitors. H1HeLa cells were either untreated/mock (M) or infected with PV at an MOI of 20 for 6 h. At 0 hpi, the infected cells were washed, followed by either adding normal DMEM (PV -NH_4_Cl) or DMEM with 20mM of ammonium chloride (PV; +NH_4_Cl), and samples were collected every hour during infection. Samples were subjected to western blot analysis for traditional autophagy markers: LC3B, p62(SQSTM1), and its Cp (cleavage product) and viral proteins: VP3 (virus structural capsid protein 3), as well as non-structural proteins, such as 2C (NSP2C) and 2B (NSP2B). Beta-actin served as a loading control.

**Figure S4.**
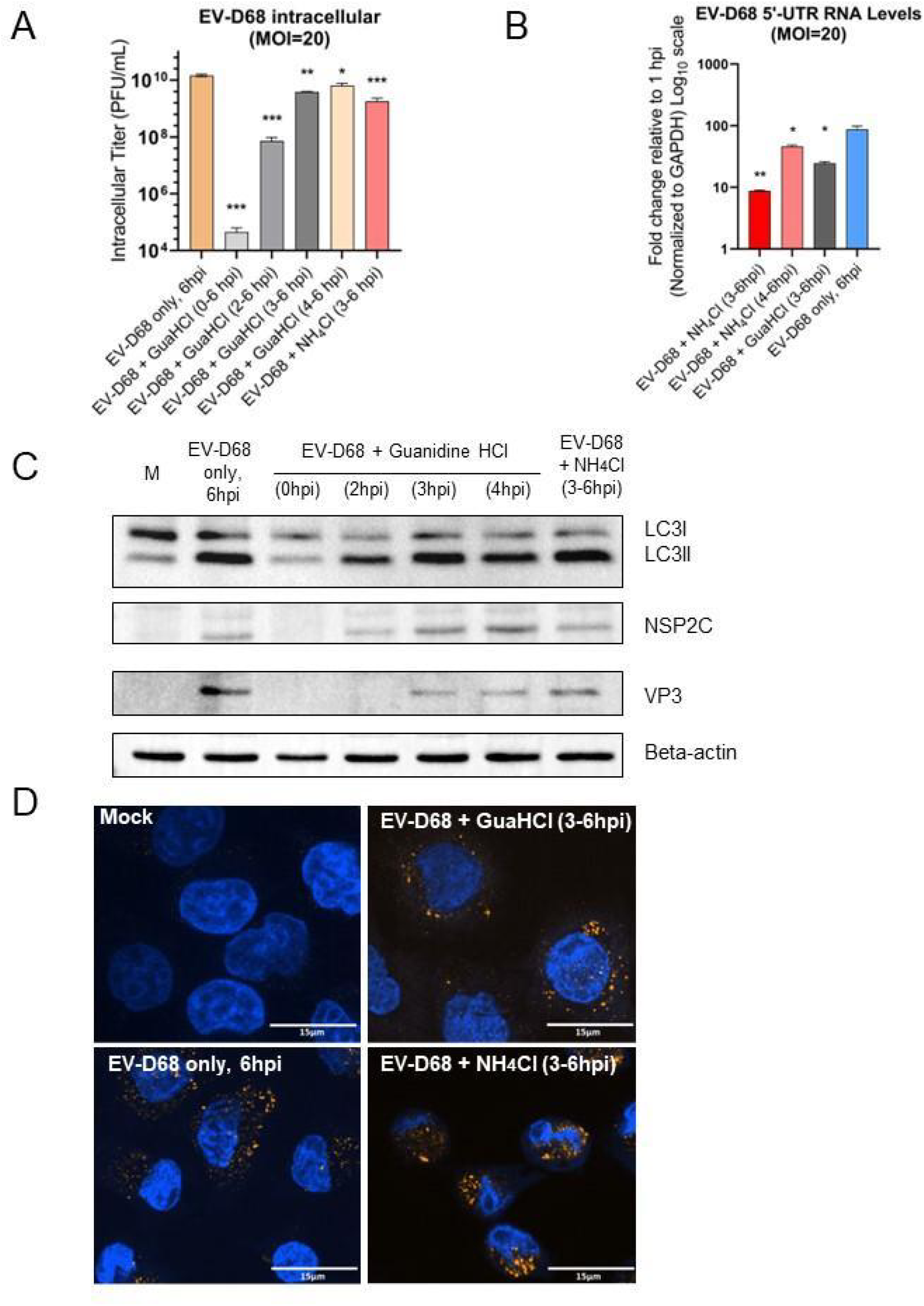
Effect of ammonium chloride and guanidine chloride treatments on EV-D68 titer, RNA levels, and protein synthesis. Related to Figure 5. (A) Titers of intracellular EV-D68 upon treatment with ammonium chloride (NH_4_Cl, 20 mM) or guanidine chloride (GuHCl, 2mM) at indicated time points. H1HeLa were infected with EV-D68 at an MOI of 20; after 30 min of virus adsorption at 37°C (0 hpi), cells were washed twice to remove the initial virus input. At the indicated time points post-infection, ammonium chloride (20mM in complete DMEM) or GuHCl (2mM) was added to the cells until the end of the infection (6hpi). Control samples (EV-D68 only) were left untreated until virus collection at 6hpi. Intracellular titers were analyzed by plaque assay. Unpaired student’s t-test was used for the statistical analysis (***= p< 0.001; **= p< 0.01; *= p ≤ 0.05; ns=not significant). (B) Effect of ammonium chloride (NH4Cl, 20 mM) and guanidine chloride (GuHCl, 2mM) on EV-D68 RNA levels assessed by qRT-PCR. Cells were infected with EV-D68 at an MOI of 20 for 30 min at 37°C, then the residual virus was washed away, followed by either adding normal media (EV-D68 only), or complete media containing 2 mM of guanidine chloride (EV-D68 + GuHCl), or 20mM of ammonium chloride at indicated time point until virus collection (6hpi). Untreated infection (EV-D68, 6hpi) served as a control. Total RNA was extracted from all samples and subjected to qRT-PCR RNA analysis following cDNA synthesis. Data represents relative expression of 5′UTR where samples are compared to the 1 hpi. All samples are normalized to GAPDH and have been log10 transformed. Unpaired student’s t-test was used for the statistical analysis (***= p< 0.001; **= p< 0.01; *= p ≤ 0.05; ns=not significant). (C) Effect of the ammonium chloride treatment on EV-D68 protein synthesis. H1HeLa cells were either untreated/mock (M) or infected with EV-D68 at an MOI of 20 for 6 h. At 0 hpi, the infected cells were washed, followed by adding normal DMEM (EV-D68 only). At indicated time points, complete DMEM with 20mM of ammonium chloride (EV-D68 + NH_4_Cl) or 2mM of guanidine chloride (EV-D68 + GuHCl), and samples were collected at 0, 2, 3, 4 and 6 hpi. Samples were subjected to western blot analysis for autophagy marker LC3B and viral proteins: VP3 (virus structural capsid protein 3), as well as non-structural protein 2C (NSP2C). Beta-actin served as a loading control. (D) Confocal imaging of EV-D68 infection upon ammonium chloride or guanidine chloride treatment added during the transition point. H1HeLa were infected with EV-D68 (MOI=20) for 6h or left uninfected (mock). Infected cells were then left untreated (EV-D68 only), treated with ammonium chloride (EV-D68 +NH_4_Cl, 3-6hpi) or guanidine chloride between 3 to 6hpi (EV-D68 + GuHCl, 3-6hpi). Cells were fixed and then stained for dsRNA (yellow) and nuclei (blue). Scale bar: 15 µM.

**Figure S5.**
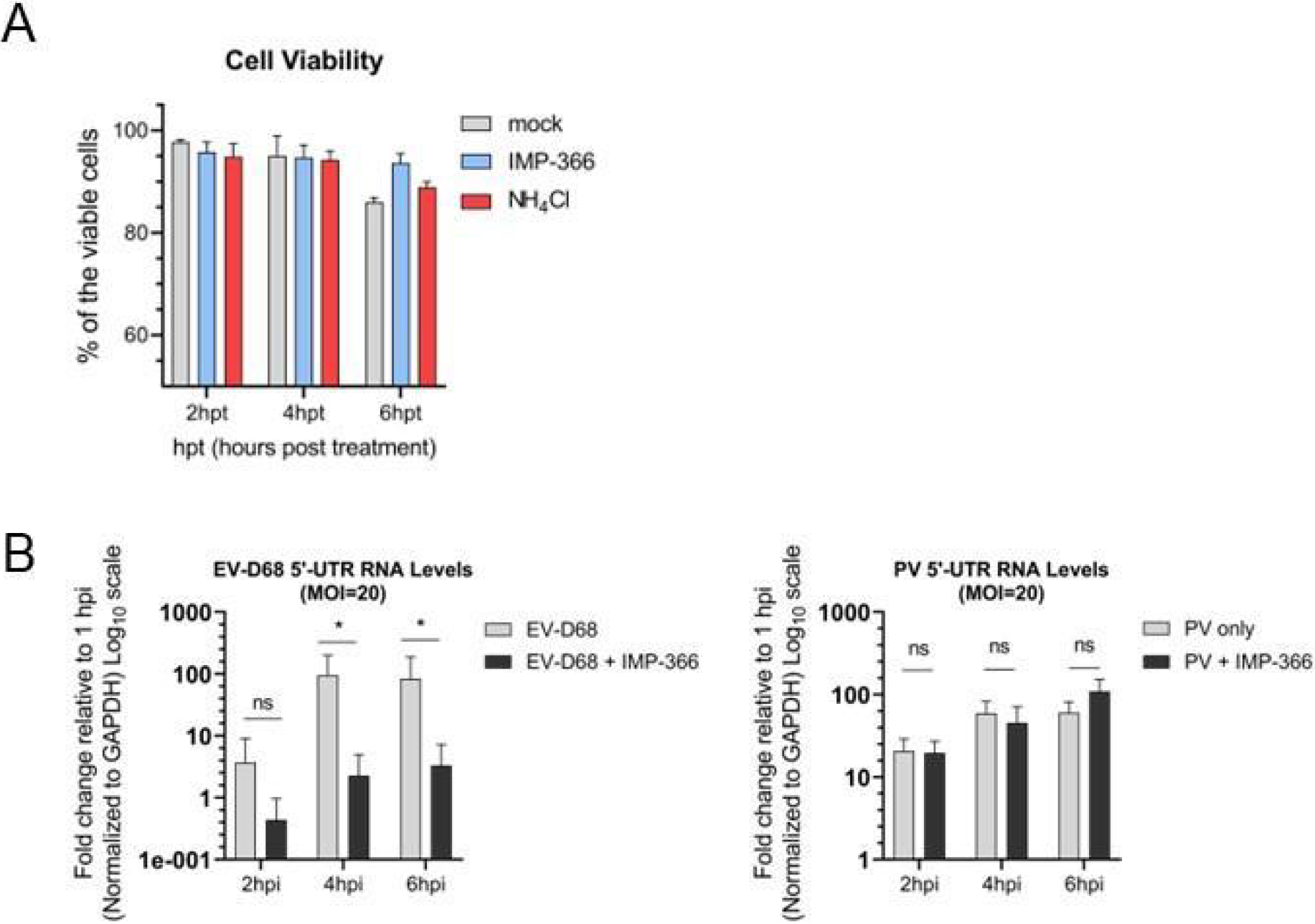
Effects of continuous treatments on H1HeLa cell viability and EV-D68 and PV RNA levels during infection. Related to Figure 6. (B) IMP-366 (5μM) and NH_4_Cl (20mM) on H1HeLa cell viability (via trypan blue assay) at the indicated times. Each data point represents the mean ± SD, n = 3. The difference in viability among treated vs. untreated cells is insignificant. (C) Effect of IMP-366 on EV-D68 and PV RNA levels assessed by qRT-PCR. Cells were infected with EV-D68 or PV at an MOI of 20 for 30 min at 37°C, then the residual virus was washed away, followed by either adding normal media (EV-D68/PV only) or media containing 5μM of IMP-366 (EV-D68/PV; +IMP-366) at 0 hpi until virus collection (6hpi). Untreated infection (EV-D68/PV) served as a control. At indicated time points, cells were collected, total RNA was extracted from all samples and subjected to qRT-PCR RNA analysis following cDNA synthesis. Data represents relative expression of 5′UTR where samples are compared to the 1 hpi. All samples are normalized to GAPDH and have been log10 transformed. Unpaired student’s t-test was used for the statistical analysis (***= p< 0.001; **= p< 0.01; *= p ≤ 0.05; ns=not significant).

